# Cohesin in space and time: architecture and oligomerization *in vivo*

**DOI:** 10.1101/2020.07.31.229682

**Authors:** Siheng Xiang, Douglas Koshland

## Abstract

Cohesin helps mediate sister chromatid cohesion, chromosome condensation, DNA repair and transcription regulation. Cohesin can tether two regions of DNA and can also extrude DNA loops. We interrogated cohesin architecture, oligomerization state and function of cohesin oligomers *in vivo* through proximity-dependent labeling of cohesin domains. Our results suggest that the hinge and head domains of cohesin both bind DNA, and that cohesin coiled coils bend, bringing the head and hinge together to form a butterfly conformation. Our data also suggest that cohesin efficiently oligomerizes on and off DNA. The levels of oligomers and their distribution on chromosomes are cell cycle regulated. Cohesin oligomerization is blocked by mutations in distinct domains of Smc3p and Mcd1p, or depletion of Pds5p. This unusual subset of mutations specifically blocks the maintenance of cohesion and condensation, suggesting that cohesin oligomerization plays a critical role in these biological functions.

## Introduction

Chromosome segregation, DNA damage repair, and the regulation of gene expression require the pairing or folding of chromosomes (Onn et al., 2008; Uhlmann, 2016). Remarkably, these different types of chromosome organizations are all mediated by a conserved family of protein complexes called SMC (Structural Maintenance of Chromosomes) (Hassler et al., 2018; Hirano, 2016; Nolivos and Sherratt, 2014; Onn et al., 2008). SMC complexes pair and fold chromosomes by two activities. First, they can tether together two regions of DNA, either within a single chromosome or between chromosomes (Hassler et al., 2018; Onn et al., 2008). Second, by combining this tethering activity with their ability to translocate along DNA, SMC complexes can also extrude DNA loops (loop extrusion) *in vivo* and *in vitro* (Davidson et al., 2019; Ganji et al., 2018; Kim et al., 2019; Wang et al., 2017). The molecular mechanisms for these activities and their regulation are still being elucidated.

SMC complexes have a conserved architecture (Hassler et al., 2018). At their core, all SMC complexes have a dimer of two evolutionarily related proteins, which are called Smc (Hirano and Mitchison, 1994; Strunnikov et al., 1995, 1993). Each Smc subunit has two globular domains, a head domain and a hinge domain, separated by a long 50 nm coiled coil. Smc subunits dimerize by two distinct interactions, one between the two heads and another between the two hinges. The separation of head and hinge dimers by their intervening long coiled coils allows the Smc dimers to achieve multiple conformations *in vitro*, including a long rod, a large ring, or a butterfly (where the coiled coils of the SMC subunits adopt a bent conformation, allowing the hinge and head domains to interact). (Bürmann et al., 2019; Hirano et al., 2001; Soh et al., 2015). The Smc subunits also act as scaffolds to bind non-Smc proteins (Yatskevich et al., 2019). The presence of different complex architectures *in vitro* raises many questions. Which, if any, of these different conformations exist *in vivo*? How do any of these conformations contribute to different SMC functions? Are there other structural features of SMC complexes, like oligomerization (dimers or multimers)? If oligomers exist, what is their function, and what factors control their formation?

One SMC complex, called cohesin, was originally discovered in budding yeast because it mediates cohesion between sister chromatids (Guacci et al., 1997; Michaelis et al., 1997). The subunits of budding yeast cohesin (Figure 1A. Smc1p and Smc3p, and the non-SMC subunits, Mcd1p/Scc1p and Scc3p) are conserved throughout eukaryotes (Guacci et al., 1997; Michaelis et al., 1997; Yatskevich et al., 2019). Sister chromatid cohesion is essential for chromosome segregation in mitosis and meiosis (Guacci et al., 1997; Klein et al., 1999; Michaelis et al., 1997). Subsequently, cohesin was also shown to be important for chromosome condensation, efficient DNA repair, and the regulation of gene expression (Bloom et al., 2018; Dorsett and Ström, 2012; Guacci et al., 1997; Lopez-Serra et al., 2013; Ström et al., 2004; Wood et al., 2010; Zhang et al., 2019). Mutations in cohesin subunits are thought to drive cancer, and cause age-dependent congenital disabilities and developmental disorders (Brooker and Berkowitz, 2014; Chiang et al., 2010; Ogawa, 2019; Romero-Pérez et al., 2019; Watrin et al., 2016).

**Figure 1.**
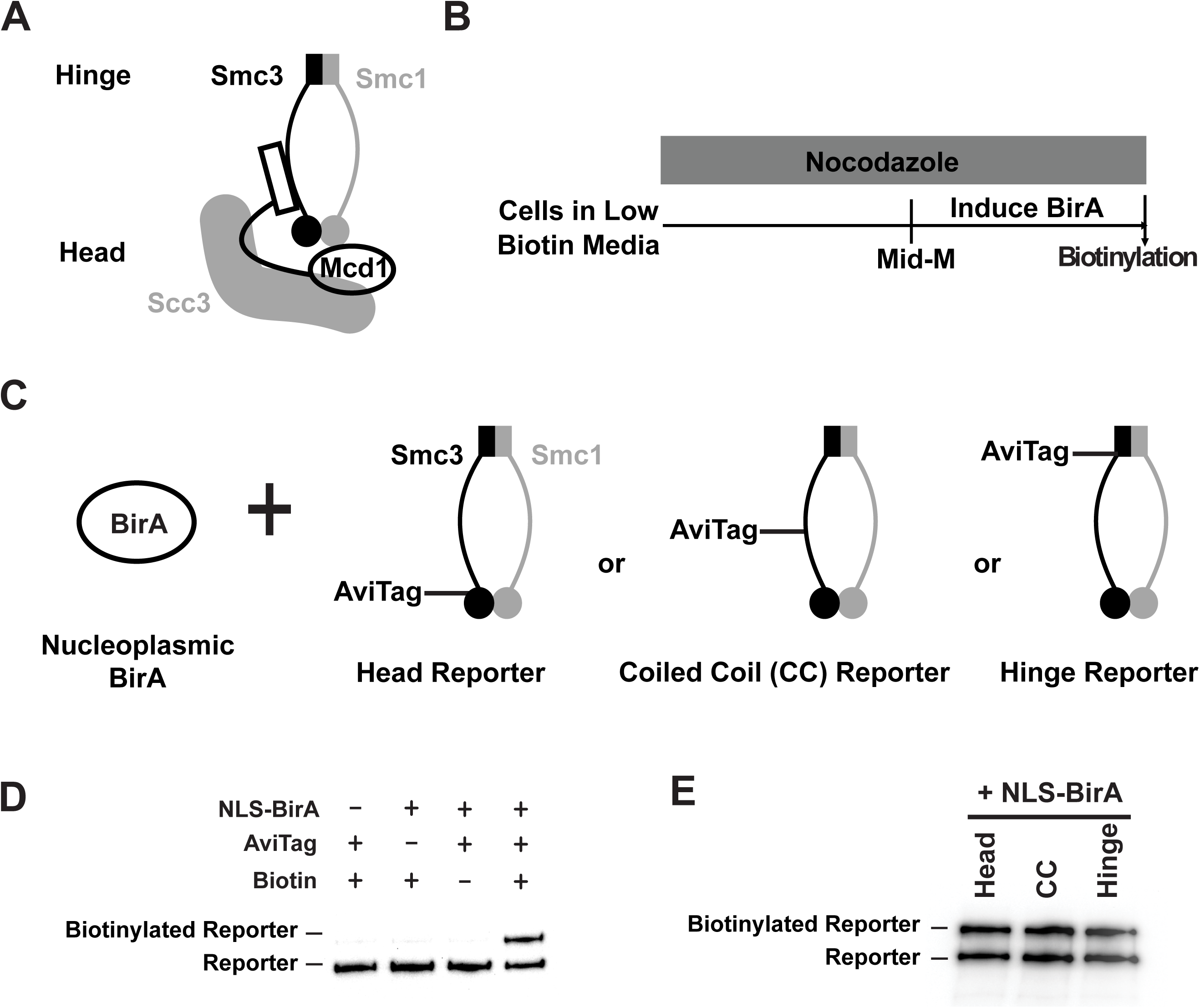
Smc3p reporters for proximity biotinylation experiments. (A) Budding yeast cohesin consists of one single copy of each of the four subunits: Smc1p, Smc3p, Mcd1p and Scc3p. (B) Experimental regimen used to test the AviTag accessibility of Smc3 reporters. Cultures of strains with *SMC3* reporter alleles were grown in low biotin synthetic media at 30 °C overnight and treated with nocodazole for 2.5 hours to arrest cells in mid-M phase. Then NLS-BirA, a nuclear localized version of BirA enzyme, was induced for one hour. 10 nM biotin was added to initiate biotinylation. After a 7-minute biotin pulse the reactions were terminated by trichloroacetic acid (TCA). (C) Cartoons depicting strains used. A nucleoplasmic BirA (NLS-BirA) was overexpressed in strains carrying one of the three *SMC3* reporter alleles. Each reporter had an AviTag fused to a flexible linker of 6xHA tag inserted into the head, the coiled coil (CC) or the hinge domains. Mcd1p and Scc3p are omitted from the cartoon for clarity. (D) Streptavidin gel shift assays to detect biotinylation of the head reporter. Cell lysates were prepared in SDS sample buffer, incubated with 1 mg/ml streptavidin for 15 minutes and subjected to Western Blot analysis using anti-HA antibodies to detect the AviTag reporter. Biotinylation of the head reporter is seen as a slower migrating band (lane 4). No reporter biotinylation was detected in cells missing BirA enzyme (lane 1), in cells with a dead AviTag (lane 2, no lysine in AviTag) or cells treated with TCA before biotinylation (lane3). (E) Comparison of AviTag accessibility to BirA mediated biotinylation. Cells were arrested in mid-M and treated with biotin pulse. Head, CC and hinge reporters were biotinylated to the same extent by a nucleoplasmic BirA.

To promote proper chromosome segregation, cells must establish cohesion in S phase. Cohesion establishment requires two steps. First, the Scc2p/Scc4p complex loads cohesin on one sister chromatid (Ciosk et al., 2000). Then cohesin is acetylated by Eco1p, which promotes tethering of the second sister chromatid (Çamdere et al., 2015; Guacci and Koshland, 2012; Rolef Ben-Shahar et al., 2008; Skibbens et al., 1999; Tóth et al., 1999; Unal et al., 2008).

Proper chromosome segregation also requires that cohesion be maintained from S phase to M phase (Hartman et al., 2000). Cohesin acetylation helps to maintain cohesion by preventing Wpl1p from removing cohesin from chromosomes (Kueng et al., 2006; Lopez-Serra et al., 2013; Rolef Ben-Shahar et al., 2008). Cohesion maintenance is also promoted by the cohesin binding protein called Pds5p (Hartman et al., 2000; Noble et al., 2006; Tanaka et al., 2001; Wang et al., 2002). Pds5p facilitates efficient Eco1p acetylation of cohesin, thereby enhancing Eco1p-dependent inhibition of Wpl1p (Chan et al., 2013). In addition, Pds5p maintains cohesion by a second unknown mechanism that is independent of Wpl1p inhibition (Tanaka et al., 2001).

The complex activities and regulation of cohesin likely depend on the presence and modulation of one or more of its conformations. The ring, rod and butterfly conformations of cohesin have been interrogated both *in vitro* and *in vivo*. Cohesin rings have been observed by electron microscopy (EM) and liquid atomic force microscopy (AFM) (Anderson et al., 2002; Hirano et al., 2001; Ryu et al., 2019). Rings have also been inferred to exist *in vivo* from experiments showing cohesin’s ability to topological entrap DNA (Haering et al., 2008). Rods have been seen by EM and have been inferred to exist *in vivo* from crosslinking experiments (Anderson et al., 2002; Soh et al., 2015); however, rods were not observed in liquid AFM images (Ryu et al., 2019). The butterfly structure is extremely rare in negatively stained electron microscopies but prevalent in liquid AFM images (Ryu et al., 2019). The existence of the butterfly structure *in vitro* is also supported by co-immunoprecipitation of recombinantly expressed hinge dimers and Scc3p. Scc3p had been shown to bind to the head domain of the native complex (Murayama and Uhlmann, 2015). *In vivo*, the existence of this structure has been inferred from physical and functional interactions of the hinge and the head-bound Pds5p (Eng et al., 2015; Mc Intyre et al., 2007). Therefore, evidence for all three cohesin conformations exist either *in vitro* or *in vivo*, but their physiological significance and relative abundances *in vivo* are unclear.

Like cohesin conformation, cohesin oligomerization has also been interrogated *in vitro* and *in vivo*. In two recent single molecular studies, the photo bleaching of fluorescently tagged cohesin came to different conclusions about whether cohesin bound to DNA as monomers or dimers (Davidson et al., 2019; Kim et al., 2019). *In vivo*, the presence of dimers in cells was tested by co-immunoprecipitation. Other studies also disagreed about the presence of dimers (Haering et al., 2002; Zhang et al., 2008). More recent *in vivo* studies revealed the functional complementation between distinct cohesin complexes (Eng et al., 2015; Srinivasan et al., 2018). This complementation could be best explained by the oligomerization of cohesins. These discrepancies make physiological significance and the relative abundance of cohesin oligomers unclear.

To interrogate cohesin conformation and oligomerization in living budding yeast, we exploited the method of proximity-dependent biotinylation (Fernández-Suárez et al., 2008; Jan et al., 2014). Our in vivo studies reveal that the Smc3p head and hinge domains are in close proximity to each other, consistent with the formation of the butterfly conformation. The head and hinge domains are in close proximity to the chromosomal DNA, suggesting that head and hinge domains bind DNA. Furthermore, our results support the formation of cohesin butterflies and cohesin oligomers, both off and on DNA. We find the abundance of cohesin oligomers and their distribution on chromosomes are regulated during the cell cycle. Finally, we show that the maintenance of cohesin oligomers requires Pds5p, the Smc3p hinge, and two regions of the Mcd1p. These results provide important new insights into the mechanism and regulation of cohesion and loop extrusion.

## Results

### Developing a proximity biotinylation method to assess cohesin structure and oligomerization *in vivo*

Proximity biotinylation is a method that exploits the ability of the bacterial enzyme, BirA, to recognize a 15 amino acid AviTag and add a biotin moiety to a lysine residue within the AviTag (Branon et al., 2018; Fernández-Suárez et al., 2008). *In vivo*, biotinylation of the AviTag is enhanced when the AviTag and BirA are fused to two interacting partners that place the BirA and AviTag in proximity. The development of proximity biotinylation to assess cohesin architecture and oligomerization was driven by three considerations. First, given that cohesin is a large complex, the level of biotinylation of the AviTag cloned into different domains of cohesin might reflect their relative proximity to a BirA partner. We constructed three alleles of the *SMC3* subunit of cohesin, one with the AviTag in the head domain (in the flexible loop after residue A1089), a second in the coiled coil (CC, after residue V966) and a third in the hinge domain (after residue P533) (Figure 1C). Strains bearing any of these AviTag-*SMC3* as sole source grew as well as wild-type strains indicating they are functional (Figure S1).

Second, we needed an assay that uniquely detects BirA-dependent biotinylation of the Smc3p reporters *in vivo*. Our assay entailed growing cells in media lacking biotin, then inducing the BirA enzyme, followed by addition of biotin to initiate biotinylation (Figure 1B). After a few minutes, we terminated biotinylation by adding TCA. The purpose of this short pulse of biotin was dual. It allowed us to limit biotinylation to specific times in the cell cycle to test temporal regulation of the interactions between our reporters and a BirA tagged partner. The biotin pulse also suppressed proximity-independent biotinylation of the reporters that occurs with extensive incubation with biotin. The biotinylated Smc3p reporter was detected as a slower mobility species due to its binding streptavidin in the protein sample buffer (Figure 1D). No biotinylation of the Smc3p head reporter was observed before the biotin pulse, in cells lacking BirA, or in cells expressing a reporter in which the critical AviTag lysine was mutated (Figure 1D). Thus, the gel shift of the Smc3p reporter was dependent upon the presence of the AviTag, BirA and biotin, validating it as a readout of BirA-dependent biotinylation of our Smc3p reporters.

Finally, we planned to use differences in the biotinylation levels of the three Smc3p-AviTag reporters to BirA tagged partners to assess cohesin conformation and cohesin oligomerization. Therefore, we needed to know whether any differences in biotinylation of the reporters could be ascribed to the AviTag accessibility to BirA in general, independent of differences in proximity to the BirA partner. To address this issue, we examined the biotinylation level of the three reporters by using a freely diffusible nucleoplasmic BirA. As shown in Figure 1E, all three reporters were biotinylated to the same extent, indicating that the head, CC, and hinge reporters are equally accessible to the unconstrained BirA enzyme. Thus, differences in reporter biotinylation likely reflect their proximity to the particular BirA fusion protein.

### Cohesin adopts a butterfly conformation *in vivo*

To test which domain(s) of Smc3p were proximal to DNA, we constructed strains bearing one of our Smc3p-AviTag reporters along with the BirA enzyme fused to a non-specific DNA binding protein from bacteria called HUα (Rouvière-Yaniv and Gros, 1975) (Figure 2A). We hypothesized that BirA enzymes tethered to DNA would only biotinylate the AviTags in the Smc3p domain(s) proximal to DNA. If cohesin bound DNA with its head domain, the AviTag reporter in the head domain would exhibit more biotinylation than AviTags in either the hinge domain or the CC. Similarly, if cohesin bound DNA with its hinge domain, the hinge reporter would be preferentially biotinylated. If DNA was embraced by cohesin, such that it was freely diffusible within the lumen formed by dimerization of the two Smc subunits, the AviTags in the head, CC, and hinge reporters should be equally biotinylated. Finally, if cohesin bound to DNA in the butterfly conformation, both head and hinge reporters would be biotinylated as both would be in proximity to DNA. In contrast, the CC reporter would be biotinylated to a lesser extent since the CC is further away from DNA.

**Figure 2.**
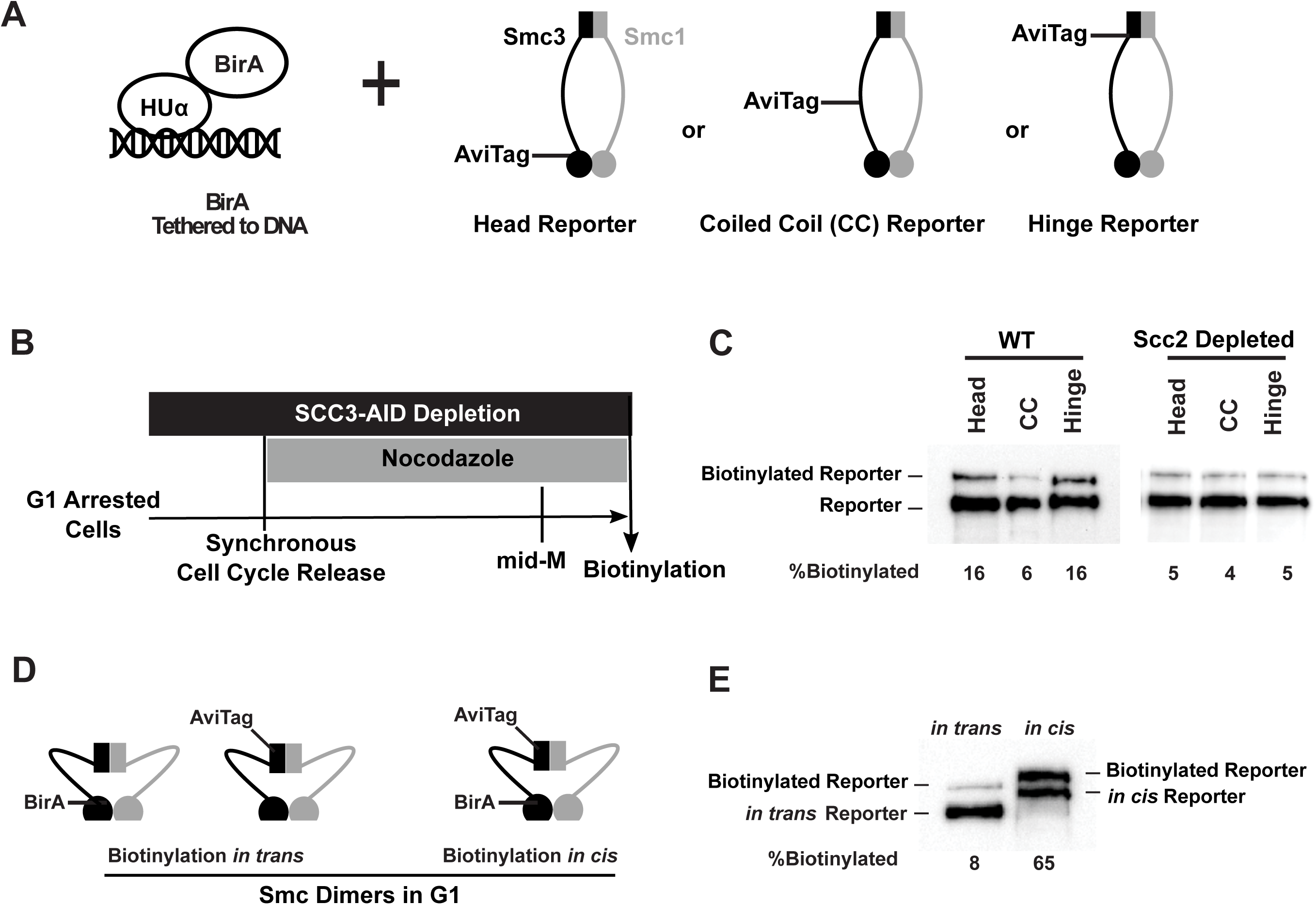
Cohesin binds DNA in a butterfly conformation with head and hinge domains on DNA. (A) Cartoons depicting strains used in this figure. BirA enzyme was tethered to DNA by HUα, a bacterial non-specific DNA binding protein. This DNA-tethered BirA was expressed in cells carrying a head, CC or hinge reporter Smc3, respectively. Mcd1p and Scc3p are omitted from the cartoon for clarity. (B) Experimental regimen used to test cohesin architecture on DNA. Cells were cultured in synthetic media with low biotin overnight to early log phase and arrested in G1 by α-factor. 1 mM auxin was added to the arrested cells to deplete the cohesin loader in *SCC2-AID* cells. Cells were synchronously released from G1 and arrested in mid-M phase in low biotin media containing auxin and nocodazole, then the cultures were treated with biotin pulse. Half of the cells were analyzed by western blotting to assess biotinylation efficiencies of the Smc3 reporters. The other half of the cultures were fixed to assay cohesin on DNA by ChIP (Figure S2). (C) Biotinylation of Smc3 reporters by DNA-tethered BirA. The gel on the left side shows biotinylation of Smc3 reporters in wild type cells. The gel on the right side shows biotinylation of Smc3 reporters in cells depleted of loader subunit *SCC2*. (D) Cartoons depicting inter- and intra-molecular biotinylation of Smc3 hinge reporter. Both strains were cultured to early log phase and arrested in G1, where cohesin complexes are not fully assembled. *In trans* biotinylation experiment was carried out in a strain carrying two *SMC3* alleles, one allele was tagged with BirA enzyme in the head domain, while the second allele was tagged with AviTag in its hinge. *In cis* biotinylation was carried out in the cells expressing double tagged Smc3, with BirA in the head and AviTag inserted in the hinge domain of the same molecule. Mcd1p and Scc3p are omitted from the cartoon for clarity. (E) Comparison of biotinylation levels of Smc3 reporters *in cis* and *in trans*. Streptavidin gel shift assay as described in Figure 1D was used to compare Smc3-AviTag biotinylation *in trans* and *in cis*.

To test these predictions, we assayed the biotinylation of the Smc3p reporters in mid-M phase when cohesin was bound to DNA. The different strains were cultured in low biotin media, arrested in G1, then released into media containing nocodazole. These cells synchronously progressed through S phase and then were arrested in mid-M phase. The mid-M arrested cells were treated with biotin pulse and then assessed for reporter biotinylation (Figure 2B). AviTags located in the Smc3p head or hinge domains were biotinylated to similar levels (about 16% of Smc3-AviTag proteins are biotinylated), while only 6% of the AviTag in the coiled coil of Smc3 protein was biotinylated (Figure 2C, left side). These results suggest that the head and the hinge domains are more proximal to chromosomal DNA than the coiled coil.

To test whether this biotinylation pattern was due to the interaction of the DNA-bound cohesin with DNA-bound HUα-BirA, we repeated the experiments in cells containing an auxin degron allele of cohesin loader subunit *SCC2* (*SCC2-AID*). Scc2p is essential for loading cohesin onto DNA. *SCC2-AID* cells were arrested in G1 phase, where the cohesin loader was depleted by adding auxin. Cells were synchronously released into nocodazole and auxin, allowing them to progress through S phase and then arrest in mid-M phase without Scc2p-AID (Figure 2B). Half of the arrested cultures were fixed with formaldehyde, and cohesin association with DNA was assessed by chromatin immunoprecipitation (ChIP) using an antibody specific for the endogenous cohesin subunit Mcd1p. The other half of the culture was subject to a short biotin pulse to allow HUα-BirA mediated biotinylation of the reporters. As expected, the Scc2p-AID depletion eliminated the Mcd1p ChIP signal at a representative cohesin associated region (CAR) (Figure S2A), confirming that cohesin loading onto chromosomes had been abolished. Importantly, we found that all three Smc3p reporters exhibited a similar low basal biotinylation level (Figure 2C, right side). Thus, the increased biotinylation of the head and hinge reporters by DNA-tethered BirA was dependent upon cohesin-DNA binding. These results confirmed that the head and hinge domains but not the coiled coil of Smc3p were close to DNA. This pattern of reporter biotinylation is consistent with cohesin binding to DNA in the butterfly confirmation or the hinge and head independently binding different regions of DNA.

To determine whether cohesin off of the DNA could form the butterfly conformation *in vivo*, we introduced both the AviTag and BirA into the same *SMC3* allele (*in cis*). The BirA was placed in the Smc3p head and AviTag in the Smc3p hinge (Figure 2D right side; biotinylation *in cis*). Intramolecular biotinylation of this double tagged Smc3p was carried out in G1 arrested cells. In G1 of yeast, Mcd1p is not expressed, so this partially assembled cohesin consists of just the Smc3p-Smc1p heterodimers. Previous studies had shown that these heterodimers are not bound to DNA (Tanaka et al., 1999). About 65% of Smc3-AviTag proteins were biotinylated in this intramolecular biotinylation experiment (Figure 2E). For comparison, we examined biotinylation when the hinge AviTag was in one Smc3p molecule and the BirA was in the head of a second Smc3p (Figure 2D left side; biotinylation *in trans*). Only 8% Smc3p-AviTag was biotinylated when BirA was fused to a second Smc3 protein (Figure 2E, biotinylation *in trans*). The high level of intramolecular biotinylation implies that the Smc3p hinge domain is often in proximity to the head domain of the same molecule independent of DNA binding. Thus the formation of the butterfly conformation appears to be an inherent property of cohesin’s Smc3p-Smc1p heterodimer.

### Cohesin oligomerizes *in vivo* and its oligomer levels are regulated during the cell cycle

If cohesin oligomerizes, then two domains from different cohesin complexes should be proximal to each other. To obtain biotinylation specific to cohesin oligomers, we constructed three strains bearing two differentially tagged Smc3p. All strains have an Smc3p with BirA inserted into its head domain. The second Smc3p in each strain has the AviTag reporter inserted at a different location, either the head, CC or hinge domains (Figure 3A). To probe for oligomerization during the cell cycle, cells were cultured in low biotin media arrested in G1, S or mid-M phase, and then subjected to a biotin pulse. Cell cycle arrests were confirmed by flow cytometry analysis of the DNA content (Figure S3A). BirA can only biotinylate an AviTag when the enzyme touches its substrate directly. Therefore, preferential biotinylation of any Smc3p reporter would occur in cohesin oligomers when the AviTagged domain was proximal to the BirA moiety located in the head of the second Smc3p (Smc3p-BirA). The maximum level of biotinylation of the Smc3p reporter would be 50% for an ordered oligomer (a dimer or a multimer like tubulin) where the domain of one cohesin touches the domain(s) of only one neighboring cohesin.

**Figure 3.**
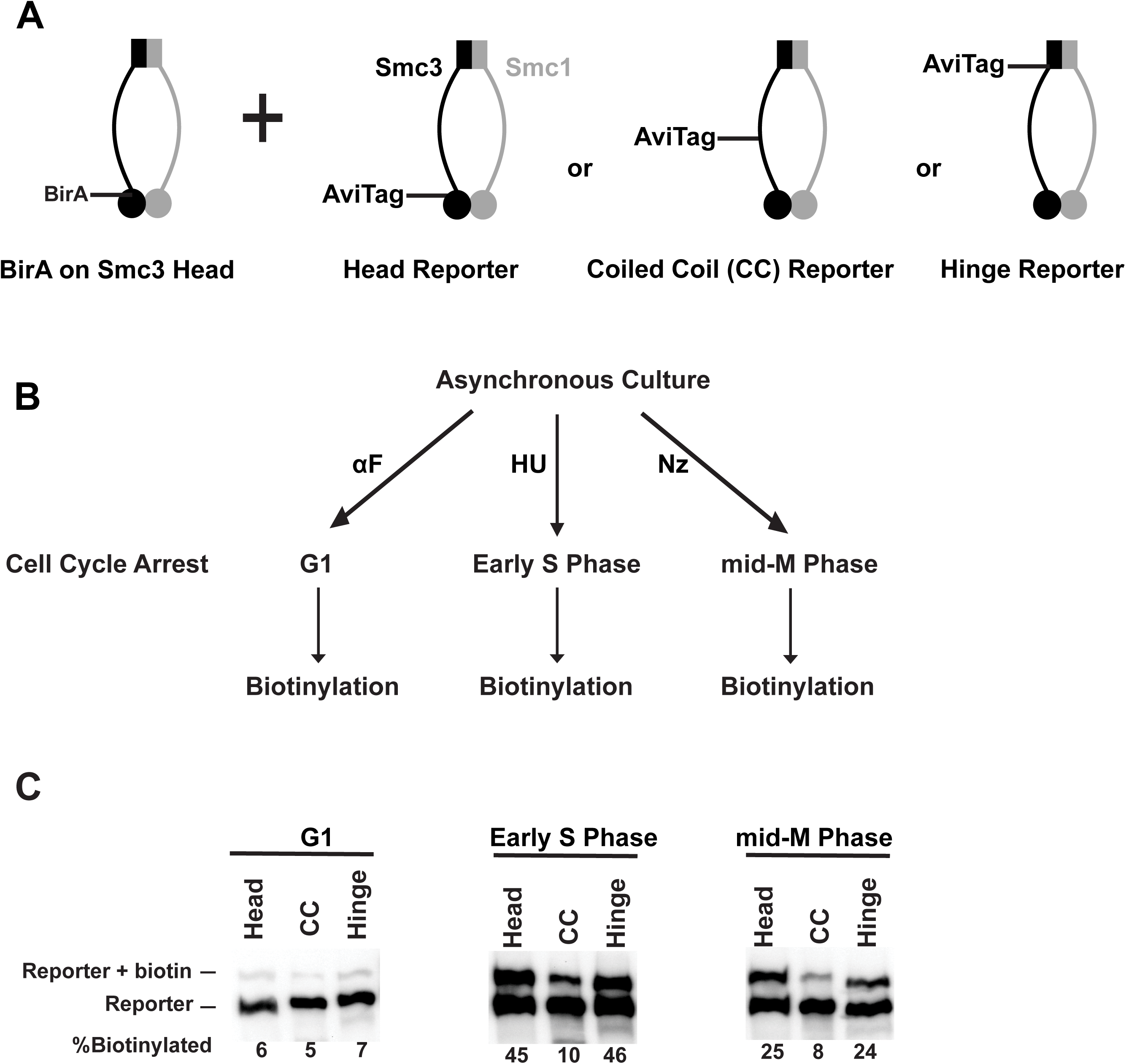
Cohesin forms oligomers that are cell cycle regulated. (A) Cartoons depicting the strains used. Each strain carries two *SMC3* alleles. One *SMC3* allele is tagged with BirA enzyme in its head domain. The other *SMC3* allele is the reporter, with AviTag fused to the head domain, the CC or the hinge domain. Mcd1p and Scc3p are omitted from the cartoon for clarity. (B) Experimental regimen to assess cohesin oligomers as a function of the cell cycle. Asynchronous cultures were treated with alpha factor, hydroxyurea or nocodazole, to arrest cells in G1, early S phase or mid-M, respectively. Cells were then treated with a biotin pulse to access levels of cohesin oligomers. (C) Comparison of oligomer formation in G1, S and mid-M. Asynchronous cultures expressing Smc3p-BirA and one of the three Smc3p reporters as shown in 3A were treated as described in 3B to assess intermolecular biotinylation of the reporters. Biotinylation was assessed by streptavidin gel shift assay as described in Figure 1D.

Biotinylation of all three Smc3p reporters were low in G1 arrested cells. Biotinylation of the CC reporter remained low in S and M phase arrested cells as well. In contrast, biotinylation of the head and hinge reporters increased 7.5 fold in S phase to 45% and 4 fold in M phase arrested cells to 25% (Figure 3C). These results suggest that cohesin oligomers form and their levels are different in cells arrested at different stages of the cell cycle.

To test whether these cell cycle differences were not unique to arrested cells but also occurred in normally dividing cells, we assayed biotinylation in cells synchronously dividing after release from G1 arrest. Cells carrying *SMC3-BirA* and the *SMC3* hinge reporter allele were cultured in low biotin media, arrested in G1, and synchronously released into a mid-M arrest. Aliquots of the culture were taken every 20 minutes, and half of the cells were treated with biotin pulse to assay Smc3p reporter biotinylation (Figure S3C). The other half of the cells from the aliquot were fixed, and DNA content in the cells was analyzed by flow cytometry. Cells exhibited a basal level of biotinylation in twenty minutes after release from G1 (Figure S3D). By forty minutes some cells started entering S phase (Figure S3E), and the biotinylation levels increased and peaked at 60-80 minutes, the latter time marked the end of S phase. Subsequently, biotinylation decreased about 50%, to a similar biotinylation level as observed in cells arrested in mid-M (Figure S3D and 3C).

These results suggest three conclusions. First, cohesins efficiently oligomerize *in vivo*. Second, oligomerization brings the head of one cohesin proximal to the head and hinge domains of another cohesin, consistent with the oligomerization of cohesins in the butterfly conformation. Third, the level of cohesin oligomers is cell cycle regulated: cohesin oligomers are absent in G1 phase, forming and peaking during S phase and are partially dissolved from G2 to M phase. The absolute levels of biotinylation suggest that most cohesin is present as oligomers in S phase and nearly half as oligomers in mid-M phase.

### Cohesin oligomers are bound to DNA and their level at CARs is cell cycle regulated

Next, we asked whether cohesin oligomers are present on chromosomes. Cohesin binds at centromeres, pericentric regions, and distinct peaks every 10-15 kb along chromosome arms called CARs (cohesin associated regions) (Blat and Kleckner, 1999; Glynn et al., 2004; Laloraya et al., 2000; Lengronne et al., 2004). We asked whether the *in trans* biotinylation of the Smc3p reporters by the Smc3p-BirA could be detected on chromosomes. We co-expressed Smc3p-BirA with a modified Smc3p head reporter in which the AviTag was placed at the end of a Myc-tag linker. This intervening linker made the biotinylated AviTag more accessible to immunoprecipitation by streptavidin beads (Figure 4A). The cells were arrested in S phase, pulsed with biotin, then fixed and processed for ChIP using streptavidin beads (Figure 4B). ChIP identified the DNA sequences associated with the Smc3p head reporter that had been biotinylated *in trans* by Smc3p-BirA (Materials and methods). DNA sequences associated with the biotinylated Smc3p marked the position of cohesin oligomers.

**Figure 4.**
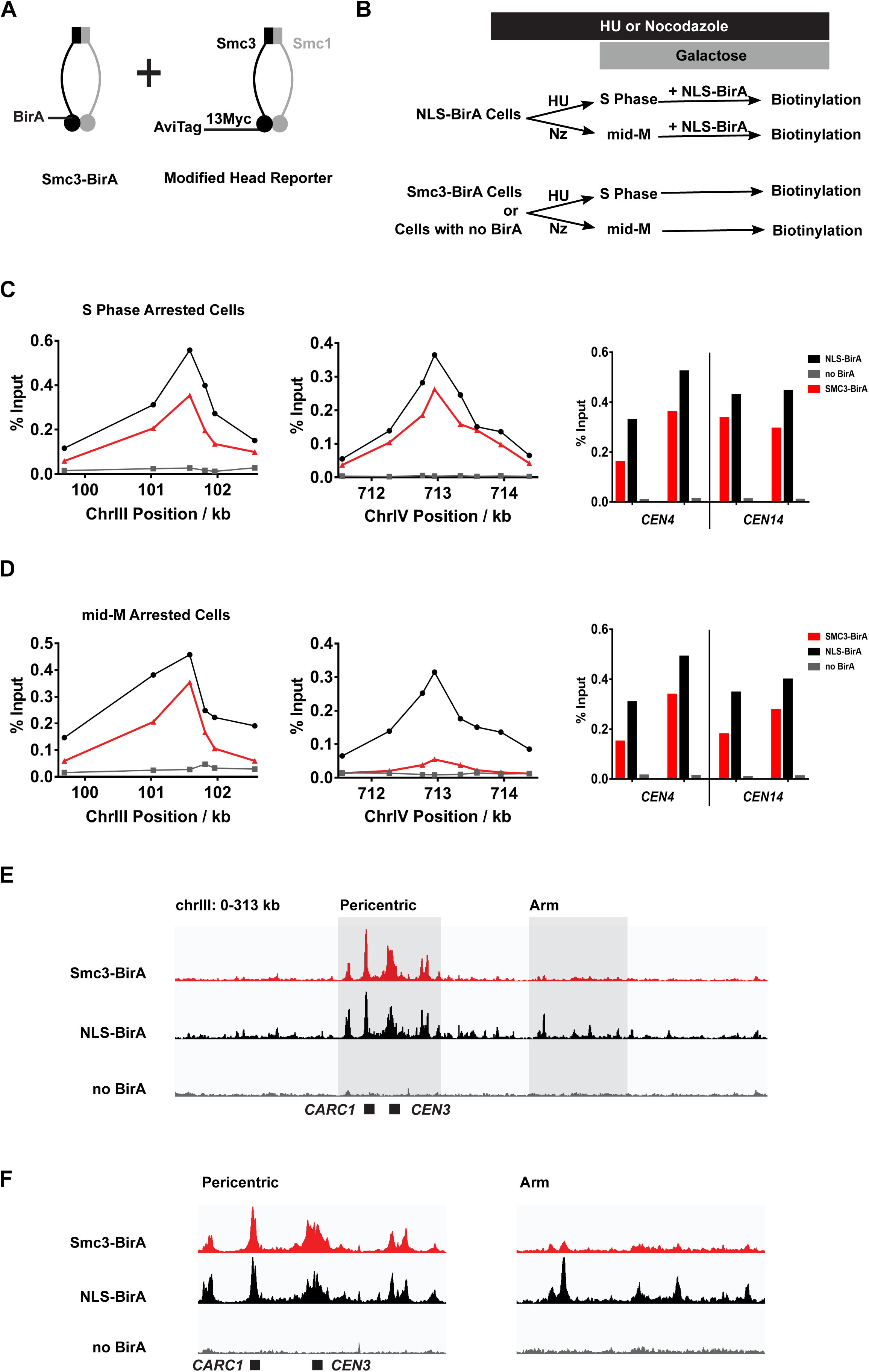
Cohesin oligomers localize to centromeres and pericentric CARs at high levels in both S and mid-M phases, but decrease at arm CARs in mid-M. (A) Strain used to assay chromosomal localization of cohesin oligomers. The strain carries two *SMC3* alleles, one expresses Smc3 with BirA in head domain (Smc3-BirA) while the other expresses modified Smc3 head reporter with an AviTag fused to Smc3 C-terminus via a 13xMyc linker (Smc3-AviTag). Mcd1p and Scc3p are omitted from the cartoon for clarity. (B) Experimental regime used to study chromosomal localization of cohesin oligomers. Cells were cultured in synthetic media, arrested in S phase or mid-M and NLS-BirA was induced in the NLS-BirA strain. Reporter biotinylation was achieved by biotin pulse and then cells were fixed and harvested for ChIP analysis as described in *Materials and Methods*. (C) Quantitative analysis of cohesin oligomer localization by qPCR in S phase arrested cells. ChIP experiments were carried out as shown in (B) with cells arrested in S phase by hydroxyurea. Cohesin oligomers in S phase arrested cells were detected on centromeres (right), pericentric CAR (left) and an arm CAR (middle). (D) Quantitative analysis of cohesin oligomer localization by qPCR in mid-M arrested cells. ChIP experiments were carried out as shown in (B) with cells arrested in mid-M by nocodazole. In mid-M arrested cells, cohesin oligomers were detected at high levels on centromeres (right) and pericentric CAR (left) but not on arm CAR (middle). (E) Chromatin immunoprecipitation (ChIP) to assess cohesin binding to chromosomes. Next generation sequencing experiments were carried out using samples in (D). Biotinylated cohesin oligomers were shown in the red trace (SMC3-BirA). Sequencing results of the control strains were plotted at the same scale. The black trace shows the sequencing result from a positive control, where all Smc3 reporter proteins can be biotinylated (NLS-BirA). The grey trace shows sequencing result from a negative control with no BirA. Positions of the centromere and a pericentric cohesin associated region (CAR) were labeled at bottom of the traces. (F) Sequencing traces zoomed in the two highlighted regions in (E). The traces on the left show the pericentric region (chrIII: 90 - 142 kb), and the traces on the right show the chromosome arm (chrIII: 178 – 237 kb).

Using qPCR we detected biotinylated Smc3p reporter at centromeres (e.g. *CEN3*), pericentric regions and at arm CARs (Figure 4C and Figure S4A). These ChIP signals were specific to reporter biotinylation as they were absent or dramatically reduced in cells lacking BirA, i.e. expressing untagged Smc3p. We wanted to assess the amount of oligomers compared with cohesin molecules present on chromosomes. Therefore, we repeated these experiments using a strain bearing the same Smc3p-AviTag reporter but over-expressing the free nuclear BirA (NLS-BirA) that we showed maximally biotinylated all the Smc3p-AviTag reporters (Figure 1E). These NLS-BirA cells exhibited a ChIP signal of biotinylated Smc3p head reporter that was similar to the *in trans* biotinylation signal at all sites examined (Figure 4C, Figure S4A). This similarity indicates that most cohesin on chromosomes are oligomers in S phase.

We wondered whether the distribution of chromosome-bound cohesin oligomers might change with the cell cycle, given the cell cycle regulated changes in oligomer abundance. We assayed for the presence of cohesin oligomers in mid-M phase arrested cells (Figure 4D and Figure S4B). As in S phase arrested cells, Smc3-BirA mediated *trans*-biotinylation of the Smc3p-AviTag reporter was robust at both *CEN* and pericentric regions. However, the *trans*-biotinylation signal was greatly reduced at some arm CARs but still above the signal in the negative control of cells lacking cells without BirA. At other CARs this signal was eliminated. At all these CARs, the biotinylation of Smc3p reporter by the free nuclear BirA was unchanged from S phase arrested cells consistent with the robust binding of cohesin to CARs in HU and mid-M arrested cells reported previously (Blat and Kleckner, 1999). These results suggest that cohesin oligomers at arm CARs are downregulated in mid-M by mechanism other than dissociating cohesin from chromosomes.

To test whether the biotinylation of these representative loci were reflective of the genome, we analyzed the mid-M sample by ChIP-seq. All peaks of cohesin oligomers correlate with previously identified regions of cohesin binding (Figure S4C). As predicted from the qPCR results, we observed robust *trans*-biotinylation signal by Smc3p-BirA at the centromeres and pericentric regions and much weaker signal or no signal at arm CARs (Figure 4E and Figure S4D). In contrast, the biotinylation signal was robust at all CARs in cells with the free nuclear BirA. The qPCR and ChIP-seq results suggest that the majority of cohesins bound to chromosomes are oligomers in S phase. In mid-M arrested cells, the presence of oligomers remains high at the centromeres and pericentric regions but are reduced on chromosome arms by mechanism independent of cohesin chromosome binding.

### Cohesin forms oligomers when fully assembled

We reasoned that the cell cycle control of cohesin assembly might contribute to the cell cycle control of cohesin oligomerization. In G1, heterodimers of Smc3p and Smc1p form, but the Mcd1p subunit is missing because *MCD1* expression is low and Mcd1p is degraded (Guacci et al., 1997). We hypothesized that the absence of oligomers in G1 (Figure 2E) may be due to the absence of Mcd1p. We introduced an *MCD1* gene under control of an inducible *GAL1* promoter into our strains with the Smc3p-BirA and the different Smc3p reporters, then compared the biotinylation of the reporters in the presence and absence of galactose (Figure 5A). In G1 arrested cells, when *MCD1* expression was repressed, a basal level of reporter biotinylation was detected (Figure 2E), but when *MCD1* was overexpressed by addition of galactose, 35% of reporters were biotinylated (Figure 5B). In mid-M arrested cells, where cohesin is already in a fully assembled complex, overexpression of *MCD1* induced by galactose did not affect cohesin oligomerization levels. Thus the absence of Mcd1p in G1 prevents cohesin oligomer formation.

**Figure 5.**
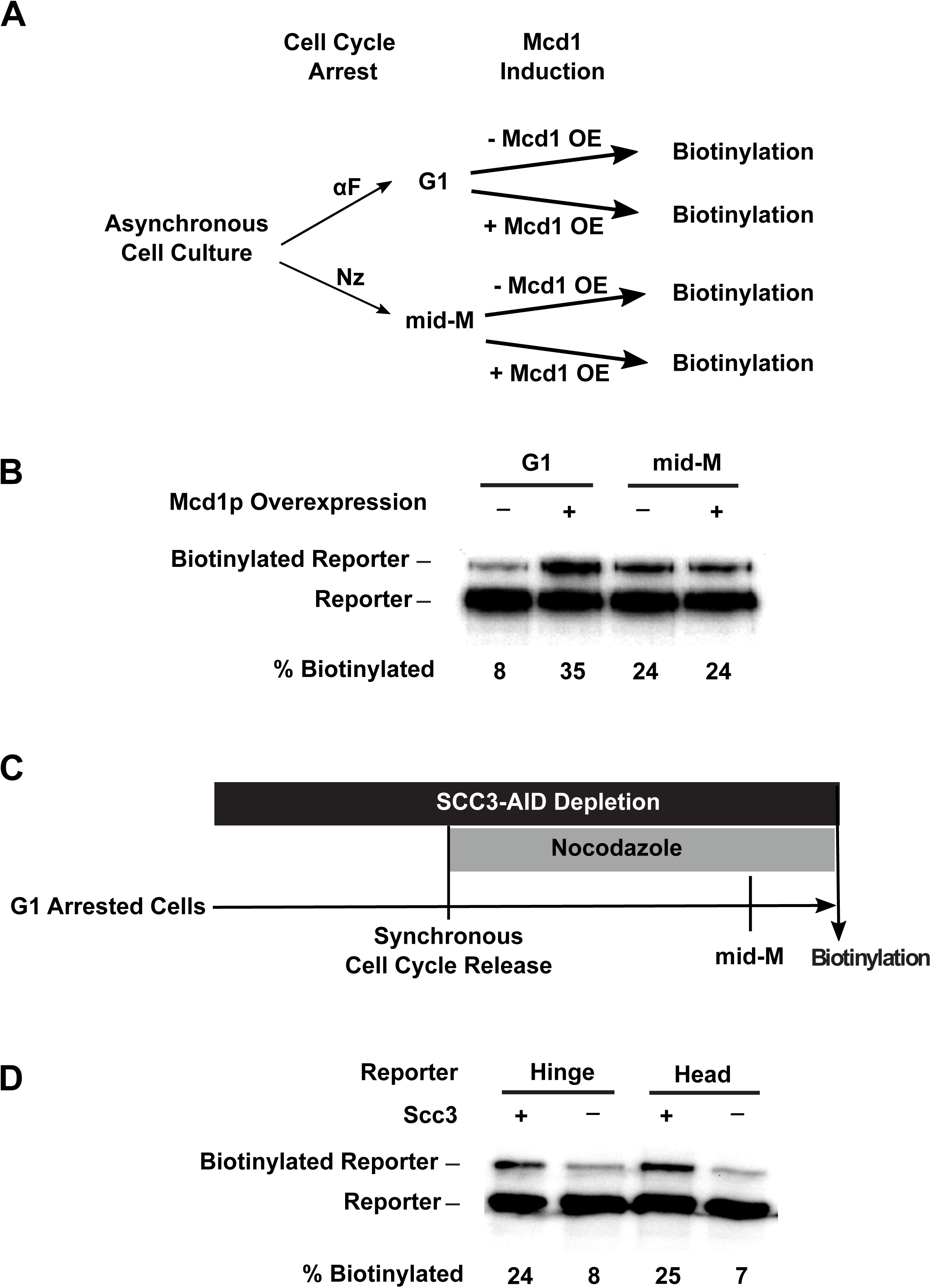
Assembly of the full cohesin tetramer is required for oligomerization. (A) Experimental regime used to assess the requirement of Mcd1p in cohesin oligomerization. The strain carries two *SMC3* alleles with one allele tagged with BirA in the head domain and the other allele tagged with AviTag in hinge domain, as shown in Figure 3A. Cultures were arrested in G1 (when Mcd1p is absent) or mid-M (when Mcd1p is assembled in cohesin tetramer). Cultures were split into half, and galactose was added to one aliquot to induce *MCD1* overexpression in arrested cells followed by a short biotin pulse to allow proximity biotinylation of the reporters. (B) Effect of Mcd1p overexpression on cohesin oligomerization. Cells arrested in G1 or mid-M phase were grown with and without Mcd1p overexpression as described in 5A. Levels of cohesin oligomers were accessed using streptavidin gel shifts as described in Figure 1D. (C) Experimental regime used to assess requirement of Scc3p in cohesin oligomerization. The strains carry two *SMC3* alleles: one allele was tagged with BirA in the head domain and the other allele tagged with AviTag in the hinge or head domains as shown in Figure 3A. Cells were cultured in low biotin synthetic media, arrested in G1, and depletion of *SCC3-AID* was carried out by auxin addition. Synchronous cell cycle release was carried out in presence of auxin and cells were again arrested in mid-M phase by nocodazole. Proximity biotinylation experiments were carried out in the mid-M phase arrested cells. (D) Scc3p is required for cohesin oligomerization. The plus signs indicate wild type cells expressing *SCC3* while the minus signs indicate cells depleted of *SCC3-AID*.

To further test the role of Mcd1p for oligomerization between G1 and M, we used an auxin degron to deplete Mcd1p (*MCD1-AID*) and assayed Smc3p biotinylation. Cells were arrested in G1, auxin added then cells were allowed to progress from G1 to mid-M arrest in the presence of auxin. This regimen prevented accumulation of newly synthesized Mcd1p-AID in S and M phases (Figure S5E). We observed only a basal level (8%) of reporter biotinylation in mid-M phase when Mcd1p was depleted, compared to 3 fold higher level of reporter biotinylation (25%) in wild type cells (Figure S5D). Thus Mcd1p absence prevents oligomerization in G1 and mid-M phases, suggesting that Mcd1p is essential for cohesin oligomerization.

Scc3p is the fourth cohesin subunit and it requires Mcd1p to assemble into cohesin (Haering et al., 2002). Therefore, Scc3p was also missing from cohesin in cells lacking Mcd1p in G1 or after Mcd1p depletion in mid-M phase cells. We tested whether Scc3p was also required for Smc3p reporter biotinylation by Smc3p-BirA. We assayed this biotinylation in cells depleted of Scc3p-AID (Figure S5B) from G1 and M using the same regimen as we used for Mcd1p-AID. We observed a reduction of biotinylation similar to the Mcd1p-AID depleted cells (Figure 5D). These results suggest that both non-Smc subunits of cohesin are required for oligomer formation.

### Pds5, but not Scc2p or Eco1p, is required for cohesin oligomerization post S phase

In budding yeast, cohesin function is modulated by four dedicated regulatory factors (Onn et al., 2008; Yatskevich et al., 2019). We assessed the potential roles of these factors in cohesin oligomerization by introducing auxin degron alleles, or deletion alleles in strains bearing Smc3p-BirA and the Smc3p-AviTag reporters. Cells that were depleted for these factors in G1 by auxin treatment or by deletion were allowed to synchronously progress to arrest in mid-M without the depleted factor. The mid-M cells were assessed for Smc3p reporter biotinylation *in trans* as a readout of oligomerization.

Scc2p is an essential subunit of the cohesin loader and is required for cohesin to bind DNA (Ciosk et al., 2000). Eco1p acetylates the Smc3p cohesin subunit, which stabilizes cohesin binding to DNA and enables establishment and maintenance of sister chromatid cohesion and condensation (Çamdere et al., 2015; Guacci and Koshland, 2012; Rolef Ben-Shahar et al., 2008; Skibbens et al., 1999; Tóth et al., 1999; Unal et al., 2008). We found that depletion of Scc2p-AID or Eco1p-AID had no effect on the *trans*-biotinylation levels seen with either the head or the hinge reporters as compared to wild type (Figures 6B & 6C). The efficacy of depletion of Scc2p-AID and Eco1p-AID was confirmed by Western (Figure S6A & Figure S6C). Moreover, cohesin failed to bind DNA in Scc2p-AID depleted cells, confirming its loss of function (Figure S6D). These results support three conclusions. Neither the loading of cohesin on DNA nor cohesin acetylation are required to form oligomers. In addition, the presence of cohesin oligomers alone is not sufficient to ensure cohesion or condensation in budding yeast.

**Figure 6.**
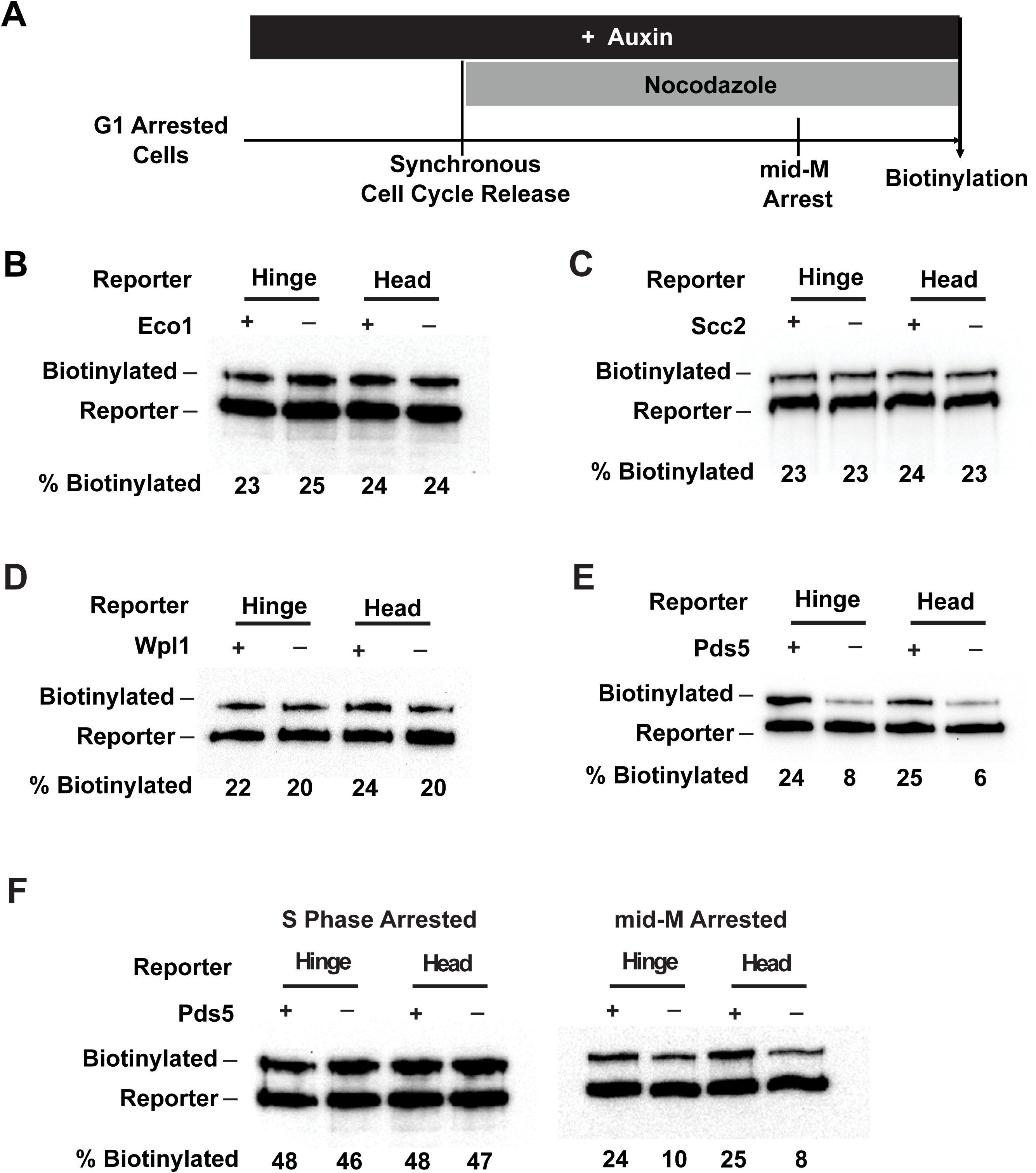
Cohesin oligomerization requires Pds5p, but not Eco1p, Wpl1p nor cohesin loader. (A) Experimental regime used to assess requirement of cohesin regulators for cohesin oligomers. Cells were cultured in low biotin synthetic media, arrested in G1, and depletion of the regulator studied was carried out by addition of auxin (except for *wpl1*Δ strains). Synchronous cell cycle release was carried out in presence of auxin and cells were again arrested in mid-M by nocodazole. Proximity biotinylation experiments were carried out in the mid-M arrested cells. (B) Eco1p is not required for cohesin oligomerization. Plus signs indicate cells carrying wild type *ECO1*, while the minus signs indicate strains depleted of *ECO1-AID*. *ECO1-AID* depletion was confirmed by western blotting (Supplementary Figure 6A). (C) Scc2p is not required for cohesin oligomerization. Plus signs indicate cells carrying wild type *SCC2*, while the minus signs indicate strains depleted of *SCC2-AID*. (Supplementary Figure 6A). *SCC2-AID* depletion was confirmed by western blotting (Supplementary Figure 6C). Cohesin is not loaded onto DNA in absence of Scc2p (Supplementary Figure 6D). (D) Wpl1p is not required for cohesin oligomerization. Plus signs indicate strains carrying the wild type *WPL1* allele, while minus signs represent *wpl1*Δ strains. (E) Cohesin failed to oligomerize in Pds5p depleted cells. Plus signs indicate cells expressing wild type *PDS5*, while minus signs indicate strains depleted of *PDS5-AID*. *PDS5-AID* depletion was confirmed by western blotting (Supplementary Figure 6B). Hinge or head reporters were biotinylated to basal level in absence of Pds5p. (F) Effects of Pds5p depletion on cohesin oligomerization in S phase or mid-M arrested cells. Cells were arrested in S phase or mid-M phase, respectively, and auxin was added to the cell cycle arrested cultures to deplete Pds5p. Cohesin oligomers were assayed by biotin pulse after Pds5p depletion. Cohesin oligomerization does not require Pds5 in S phase arrested cells but requires Pds5p in mid-M arrested cells.

Another regulatory factor called Wpl1p dissolves cohesion by removing cohesin from chromosomes but a subset of cohesin is made refractory to Wpl1p by Eco1p acetylation (Chan et al., 2013; Kueng et al., 2006). Wpl1p inhibits condensation by an unknown mechanism (Guacci and Koshland, 2012; Lopez-Serra et al., 2013). In cells harboring a deletion of *WPL1* (wpl1Δ), the biotinylation levels of the head or the hinge reporters were indistinguishable from that observed in wild type (Figure 6D). These results suggest that Wpl1p is neither an activator nor an inhibitor of oligomerization.

The fourth factor, Pds5p, is required for the maintenance of cohesion and condensation in budding yeast (Hartman et al., 2000; Noble et al., 2006; Tanaka et al., 2001; Wang et al., 2002). It is also necessary for efficient Eco1p-dependent acetylation of cohesin during S phase (Chan et al., 2013). In contrast to other regulators, Pds5p-AID depletion dramatically reduced the biotinylation levels of both the head and the hinge reporters (Figure 6E). The cohesin biotinylation level observed without Pds5p was comparable to the basal level observed in G1 arrested cells that lack Mcd1p or mid-M phase cells depleted for Mcd1p. Pds5p-AID depleted cells have reduced cohesin acetylation and DNA binding (Chan et al., 2013). However, our analysis of Scc2p-AID and Eco1p-AID depletion showed that Smc3p-reporter biotinylation was independent of either acetylation or DNA binding (Figures 6B & 6C). Therefore, Pds5p promotes cohesion oligomerization in mid-M by a mechanism distinct from its function in cohesin acetylation or cohesin binding to DNA.

DNA-independent oligomerization of cohesins suggested the possibility that Pds5p might be required for the cohesin oligomerization of DNA-free cohesin. To test this prediction, we examined Smc3p reporter biotinylation in cells depleted for the cohesin loader Scc2p-AID, Pds5p-AID, or co-depleted for Scc2p-AID and Pds5p-AID. When Scc2p and Pds5p were both depleted, the biotinylation levels of both the head or the hinge reporters was reduced to a level comparable to the Pds5p-AID depletion alone and reduced about 70% compared to Scc2p-AID depletion alone (Figure 6E and S6J). These results show that Pds5p promotes cohesin oligomerization independent of DNA binding.

In these experiments, Pds5p-AID was inactivated from G1 phase through to mid-M phase. To assess Pds5p’s role in oligomerization specifically in S and M phases, we depleted Pds5p in cells after they were arrested in S or mid-M and then assayed Smc3p reporter biotinylation while cells remained arrested. Pds5p-AID depletion in S phase arrested cells had no effect as we found the same high level of biotinylation on both the head and hinge reporters as seen in wild type cells (Figure 6F, left side). In contrast, depletion of Pds5p-AID after cells were arrested in mid-M reduced biotinylation of both reporters to basal levels (Figure 6F, right side), similar to the level upon Pds5p-AID depletion from G1 to M phase. These results suggest that cohesin oligomers can form in the absence of Pds5p, but they require Pds5p to persist post S phase. Furthermore, they further correlate cohesin oligomers with sister chromatid cohesion. Oligomers are present in S phase when cohesion is established and are maintained by Pds5p after S phase when cohesion must be maintained. We wondered whether Pds5p might be needed for the formation of the butterfly conformation since the pattern of labelling for the Smc3p reporters suggested that this conformation is present in oligomers (Figure 3C). To test this idea, we generated a Pds5-AID strain containing one *SMC3* allele bearing both BirA in its head and the AviTag in its hinge (Figure 2D, right side). We depleted Pds5p-AID in mid-M arrested cells and then subjected them to a biotin pulse. We observed robust biotinylation of the hinge AviTag (Figure S6H), indicating that Pds5p is not needed for formation of the butterfly conformation. Furthermore, it suggests that the formation of the butterfly conformation is not sufficient for oligomerization.

### Cohesin oligomerization correlates with cohesion maintenance and *RDN* condensation

Previous studies identified other cohesin mutants that are phenotypically very similar to Pds5p-depleted cells. These mutants also have defects in the maintenance of cohesion and condensation. These similarities suggested that they identify domains of cohesin subunits important for oligomerization. Therefore, we examined these mutants for cohesin oligomerization.

We first examined one such mutant, *mcd1-V137K*. The Mcd1p-V137K fails to bind to Pds5p and cannot support cell viability (Chan et al., 2013; Eng et al., 2014)). Therefore, we generated strains bearing two *MCD1* alleles, *mcd1-V137K* and *MCD1-AID* to enable viability. This strain also contained two *SMC3* alleles to enable cohesin oligomerization to be assessed, one has Smc3p-BirA in the head and the Smc3p reporter carrying AviTag in the hinge. We depleted Mcd1p-AID in G1 arrested cells then released cells from G1 into media containing auxin and nocodazole to arrest cells in mid-M phase with mcd1p-V137K as sole Mcd1p protein in the cell (Figure 7A). We then assessed for Smc3p reporter biotinylation with a biotin pulse. Similar to Pds5p-AID depleted cells, cohesin with mcd1p-V137K had a basal level of Smc3p reporter biotinylation (Figure 7B). These results suggest that Pds5p must interact with cohesin to promote oligomerization, and thus, Pds5p likely promotes oligomerization by acting directly on cohesin.

**Figure 7.**
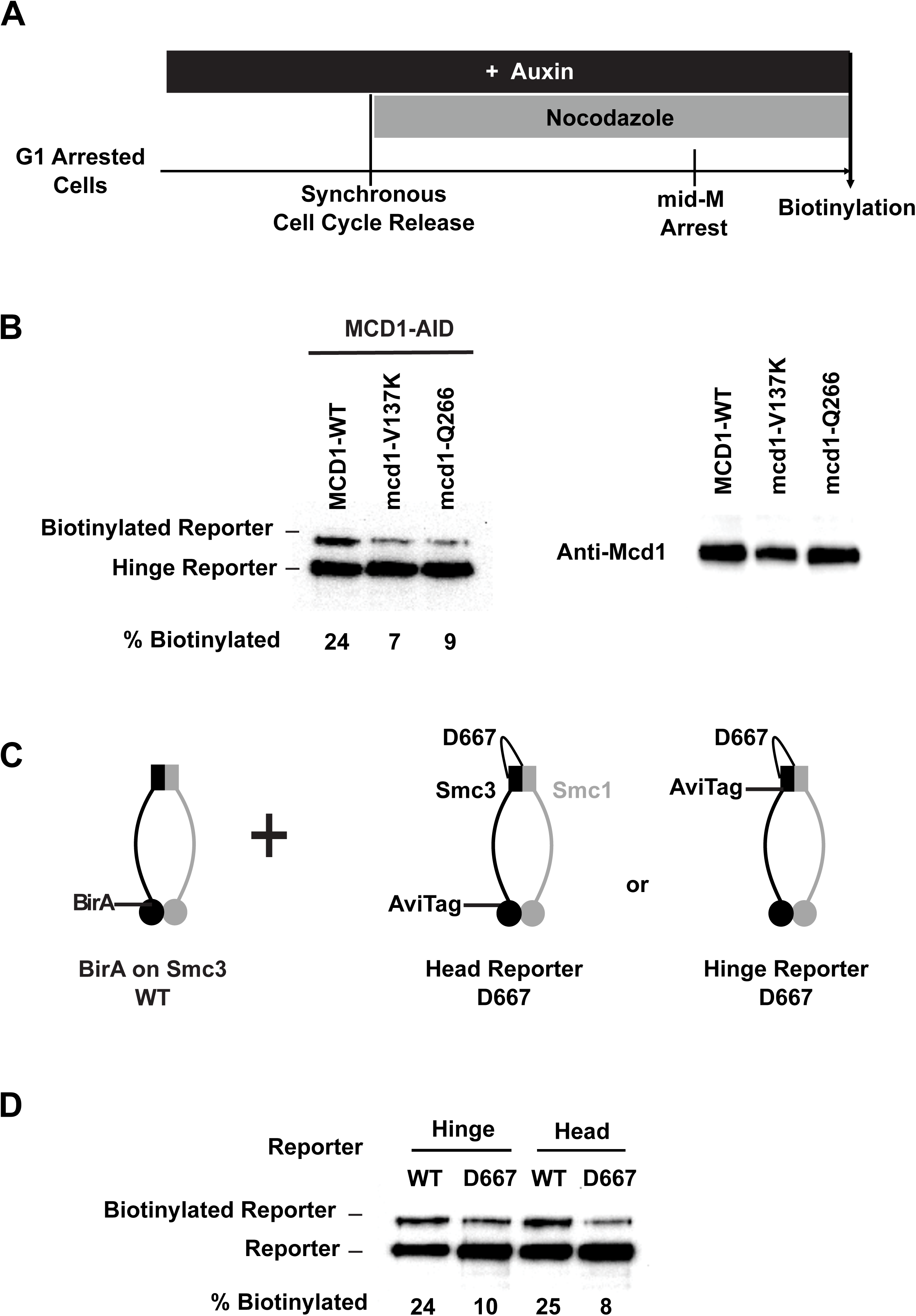
Cohesin mutants defective in cohesion maintenance failed to oligomerize. (A) Experimental regime used to assay cohesin oligomers in strains expressing wild type or mutated Mcd1p. Each strain expresses both *SMC3-BirA* and a *SMC3* reporter allele. Cells were cultured in low biotin synthetic media, arrested in G1, and depletion of wild type *MCD1-AID* was carried out by addition of auxin. Synchronous cell cycle release was carried out in presence of auxin and cells were again arrested in mid-M phase by nocodazole. Proximity biotinylation experiments were carried out in the mid-M phase arrested cells. (B) Cohesin failed to form oligomers in cells carrying either *mcd1-Q266* or *mcd1-V137K* mutation. Each stain carries two *MCD1* alleles: an auxin-depletable *MCD1-AID* and a test *MCD1* mutant allele. Lane 1 has protein extracts from cells expressing *MCD1-AID* and wild type *MCD1*; lane 2 has protein extracts from cells expressing *MCD1-AID* and *mcd1-V137K*; lane 3 has protein extracts from cells expressing *MCD1-AID* and *mcd1-Q266*. Cohesin oligomers were assayed by biotinylation (left side) and Mcd1p expression were confirmed by western blotting using Mcd1 antibody (right side) (C) Cartoons showing *SMC3* alleles used in (D). Each strain carries two *SMC3* alleles. One allele expresses a wild type, BirA tagged version of *SMC3*. The other allele expresses a wild type or *D667* mutated version of Smc3 reporter (AviTag inserted in head or hinge). Mcd1p and Scc3p are omitted from the cartoon for clarity. (D) smc3p-D667 reporters failed to oligomerize with wild type Smc3p-BirA. Cells depicted in (C) were arrested in mid-M phase and interaction between wild type Smc3p and mutated Smc3p reporters were assayed by proximity biotinylation.

This result made us wonder whether Pds5p recruitment to cohesin was sufficient to promote its oligomerization. We previously generated five amino acid insertions in Mcd1p (after residue Q266) and Smc3p (after residue D667) (Eng et al., 2014; Robison et al., 2018). The Q266 residue lies in the linker region of Mcd1p and the D667 residue is in the hinge of Smc3p. Like *mcd1-V137K*, these insertions cannot support viability and also cause defects in the maintenance of cohesion and condensation. However, cohesin containing mcd1p-Q266 or smc3p-D667 bind chromosomes and recruit Pds5p. By examining Smc3p reporter biotinylation in strains carrying these alleles, we could address whether Pds5p binding to cohesin was sufficient for cohesion oligomerization.

Using a strategy similar to our analysis of mcd1p-V137K, we generated strains that expressed only mcd1p-Q266 from G1 to mid-M arrest. We also generated strains carrying D667 mutated versions of Smc3p reporters and a second wild type Smc3p bearing BirA (Figure 7C). The *trans*-biotinylation of the head and hinge of cohesin containing mcd1p-Q266 or smc3p-D667 was reduced to levels similar to that seen in cells depleted for Pds5p-AID (Figure 7B and Figure 7D). Since Pds5p binds to cohesin bearing either mcd1p-Q266 or smc3p-D667 mutants, Pds5p binding to cohesin is necessary but not sufficient for oligomerization. Furthermore, the fact that Pds5p-depletion, smc3p-D667 and mcd1p-Q266 are all defective for oligomerization and the maintenance of cohesion and condensation, strengthen the correlation between these cohesion functions and cohesin oligomerization.

## Discussion

Here, we use a biotin-based proximity labeling to probe the *in vivo* architecture of cohesin, both on and off DNA. Our results suggest that the head and hinge domains of cohesin, but not the coiled coil are proximal to DNA. These results are consistent with DNA binding to the head and hinge but not freely floating in the coiled coil as proposed for the embrace model. Similarly, AFM imaging of another SMC complex called condensin showed DNA binding only to its head and hinge domains (Ryu et al., 2019). Moreover, the presence of two distinct DNA binding sites on cohesin was predicted from mutants that uncoupled cohesion from stable DNA binding, and from the discovery of its loop extruding activity (Davidson et al., 2019; Eng et al., 2014; Kim et al., 2019).

DNA binding to the head and hinge could occur if cohesin is in the ring, rod or butterfly conformation. We show that BirA in the Smc3p head can biotinylate an AviTag *in cis* in the Smc3p hinge. This result indicates that the head and hinge can be in close proximity within one cohesin complex *in vivo*, consistent with the presence of the butterfly conformation. This conclusion supports genetic and FRET studies that had suggested functional and physical interactions between the head and hinge domains of cohesin *in vivo* (Mc Intyre et al., 2007; Robison et al., 2018). Interestingly, the existence of the butterfly conformation off and on DNA has been observed in AFM imaging of condensin (Ryu et al., 2019). Also, super-resolution imaging of worm meiotic chromosomes has revealed the proximity of the head and the hinge domains of cohesin containing the COH-3/COH-4 (paralogs of Mcd1p) (Köhler et al., 2017). These results are consistent with the formation of a butterfly conformation on DNA. We suggest that the butterfly conformation is a conserved feature of SMC complexes *in vivo* (Figure 8B and 8C).

**Figure 8.**
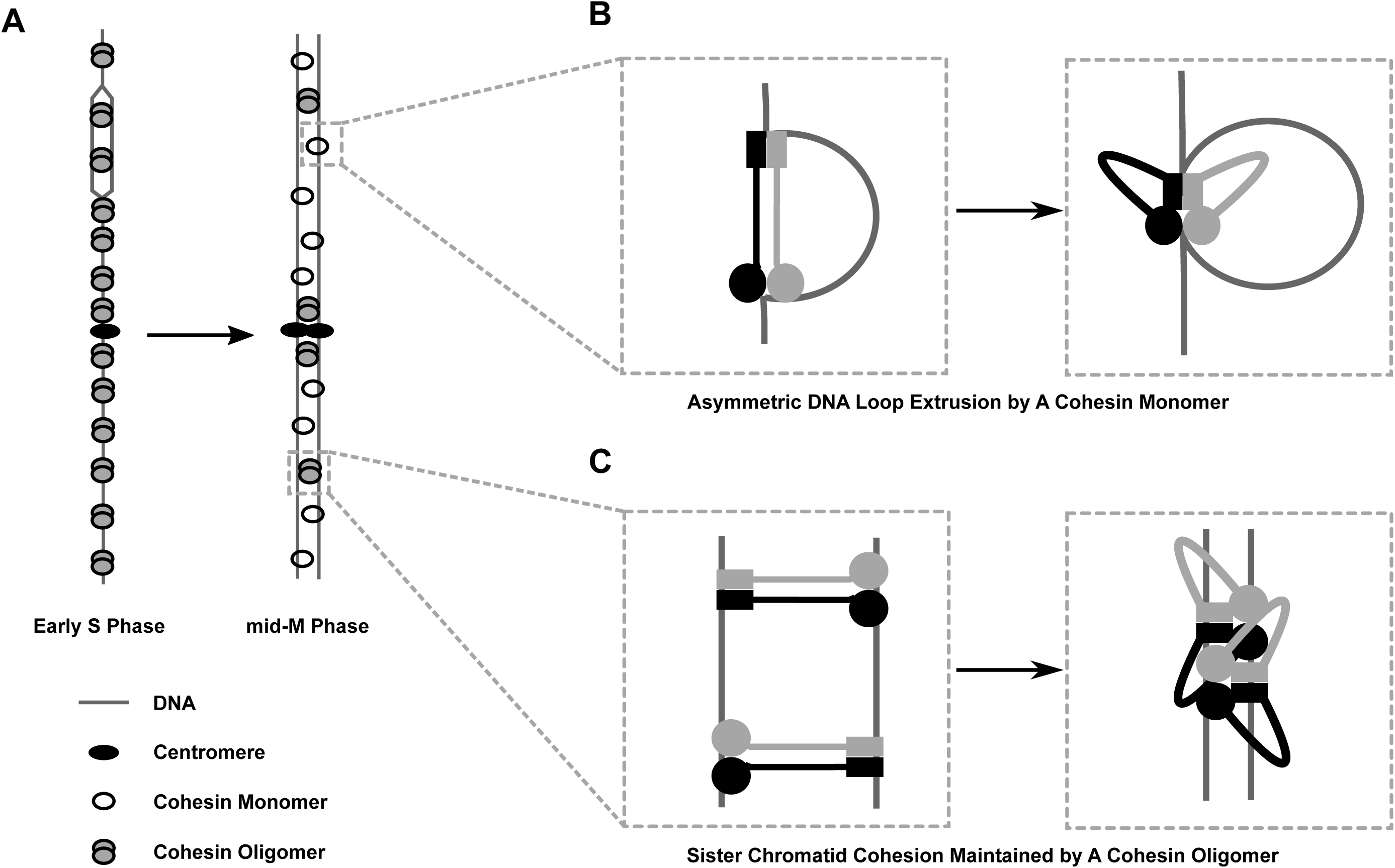
Model of cohesin oligomer regulation during cell cycle. (A) Cartoon showing regulation of cohesin oligomers during cell cycle. Cohesin complexes form oligomers at all previously identified binding sites at early S phase including centromeres, pericentric CARs and arm CAR sites. Most of the oligomers at chromosome arms dissolve in mid-M phase. A small fraction of cohesin complexes remain in the oligomeric state and keep sister chromatids tethered. Pds5 protein participates in stabilization of cohesin oligomers through S phase into mid-M. (B) Cartoon showing the initial steps of asymmetric loop extrusion by cohesin. Mcd1p, Scc3p and Pds5p are omitted from the cartoon for clarity. Cohesin monomer binds DNA with both head and hinge domains (cohesin complex in extended conformation may adopt rod or open ring structures), then the coiled coil domain bends to form the butterfly conformation and extrude a DNA loop. In subsequent steps (not shown here), DNA is transferred from the hinge to a second DNA binding site on the head. The hinge binds a new region of DNA and repeats the cycle, increasing the size of the loop. Cohesin dimmers can extrude DNA symmetrically. This model has been proposed for condensin complex (Ganji et al., 2018). (C) Cartoon showing sister chromatid cohesion by cohesin. Cohesin binds sisters with its head and hinge domains, respectively, and tethers sister chromatids. This tethering state proceeds to the butterfly conformation where it is trapped. Oligomerization of cohesins in the butterfly conformation stabilizes tethering between sisters and maintains cohesion.

The existence of the butterfly conformation, coupled with a recent scrunching model for loop extrusion by condensin, provides a working model for the mechanism of cohesion. In the scrunching model (Ryu et al., 2019), two DNA regions of the same DNA molecule are captured by the head and hinge of an extended condensin. Subsequent bending of the coiled coil to form the butterfly allows the transfer of the DNA from the hinge to the head. The coiled coil then extends, in a conformation, allowing the hinge to bind another region of DNA. Repeated cycles of scrunching cause loop extrusion (Figure 8B). We previously suggested cohesin could be converted from extruding DNA to tethering DNA by blocking the release of ADP from cohesin, thereby trapping an intermediate in the loop extrusion cycle (Çamdere et al., 2018). Here, we speculate that the hinge and head of an extended cohesin, each bind a different sister chromatid at or near the DNA replication fork (Figure 8C). Upon subsequent formation of the butterfly conformation, cohesin acetylation and possibly association with other factors prevent ADP release that is needed to dissolve the butterfly. Trapping of this intermediate prevents additional cycles of scrunching that would release the sister DNA molecule, and as a result generates a stable tether between sister chromatids.

Our biotin-based proximity labeling also suggests that cohesin forms oligomers, both off and on DNA. We showed that BirA present on one cohesin can *trans*-biotinylate the AviTag present on another cohesin when off of DNA (Scc2p depletion) or on DNA (ChIP). Oligomerization is likely driven by specific interactions between multiple cohesin domains since *trans*-biotinylation is blocked by specific mutations in Mcd1p and Smc3p (this study). Oligomerization of butterfly cohesins provides a simple explanation for why BirA at the head of one Smc3p can biotinylate *in trans* AviTags in either the head and the hinge of another Smc3p, but not its coiled coil (Figure 8C, this study). The specificity of the labeling and its occurrence off and on DNA make it unlikely to occur by other mechanisms like random aggregation or by DNA-dependent phase separation (Ryu et al., 2020).

The existence of cohesin oligomers is consistent with the *in vitro* detection of mammalian cohesin dimers, the co-immunoprecipitation of differentially tagged cohesins isolated from mammalian cells, and functional cooperation between mutated yeast cohesins *in vivo* (Eng et al., 2015; Kim et al., 2019; Srinivasan et al., 2018; Zhang et al., 2008). A recent *in vivo* imaging study of worms suggests that clusters (∼4) of cohesins containing COH-3/COH-4 bind at distinct loci on meiotic chromosomes (Woglar et al., 2020). These clusters and the proximity of the head and hinges of COH-3/COH-4 cohesins is also consistent with the oligomerization of cohesins in the butterfly conformation (Köhler et al., 2017; Woglar et al., 2020). Thus, oligomers of cohesin butterflies may be conserved in eukaryotes.

The robustness of *trans*-biotinylation suggests that a substantial fraction of cohesin is in oligomers in the cell, and therefore oligomers are likely an important feature of cohesin function (this study). Indeed, we show that *trans*-biotinylation between cohesins is blocked in four mutants (*pds5-AID, mcd1-Q266, mcd1-V137K*, and *smc3-D667*). Among alleles affecting cohesin function, these mutations have a unique phenotype: they establish but fail to maintain cohesion (Eng et al., 2014; Hartman et al., 2000; Robison et al., 2018). These results provide compelling evidence that the maintenance of oligomers is essential for maintaining cohesion.

These mutants also fail to condense the RDN locus of yeast ((Eng et al., 2014; Hartman et al., 2000; Robison et al., 2018)). This suggests a model in which cohesin oligomerization in the RDN promotes condensation. Indeed, cohesin-driven compaction, called vermicelli, has been observed in mammalian cells depleted of Wapl (Tedeschi et al., 2013). However, RDN but not vermicelli compaction requires condensin making this model unlikely (Lavoie et al., 2000; Strunnikov et al., 1995; Tedeschi et al., 2013). Alternatively, we recently showed that polo kinase binding to cohesin at the RDN is critical for proper condensin function (Lamothe et al., 2020). This recruitment may be augmented by cohesin oligomers.

In a previous study, we suggested two potential roles of cohesin oligomers in cohesion (Eng et al., 2015). Cohesion could be achieved by oligomerization of two cohesins, each bound to a sister chromatid (handcuff model). In this model, oligomers are required for cohesion establishment and maintenance. Alternatively, a single cohesin could have two DNA binding sites, and therefore be competent to form cohesion. Lateral oligomerization of individually cohesion-competent cohesins would generate sites of multivalent cohesion. In this second model, oligomerization would not be obligatory for cohesion establishment. Rather, the multivalent cohesion of the oligomer would ensure that cohesion persists at a specific locus even when tethering of individual cohesins is disrupted. We prefer the second model since *in vivo* and *in vitro* studies suggest that individual cohesins have multiple DNA binding sites (this study and (Ryu et al., 2019)), and a single cohesin is competent to form cohesion (Haering et al., 2008). In addition, our mutant analysis supports a role for oligomers in cohesion maintenance. However, to eliminate a role for oligomers in cohesion establishment will require identifying additional mutations that block cohesin oligomerization in S phase.

Pds5p was known to maintain cohesion by an unknown mechanism that was independent of Wpl1p and Eco1p (Eso1p) (Tanaka et al., 2001). This unknown mechanism likely involves Pds5p’s promotion of cohesin oligomerization given the correlation between oligomer and cohesion maintenance, and Pds5p’s ability to stabilize oligomers independent of Eco1p (this study). Pds5p may indirectly promote oligomer maintenance by modulating a post-translational modification that alters oligomer formation or dissolution. If so, this modification is not cohesin acetylation since Eco1p is not needed for oligomerization (this study). Alternatively, Pds5p may act directly to stabilize oligomers by binding to different regions of two adjacent cohesin complexes within the oligomer. This model predicts that mutations in one of these two Pds5 binding regions would block the formation of cohesin oligomers but would have no effect on Pds5p binding to cohesin. Indeed, *mcd1-Q266* and *smc3-D667* mutant cohesin still bind Pds5p but prevent oligomer formation (this study) (Eng et al., 2014; Robison et al., 2018).

The presence of cohesin oligomers *in vivo* and their abrogation in our mutants also inform on loop extrusion in budding yeast. We recently showed that cohesin forms defined loops genome-wide with CARs as loop anchors (Costantino et al., 2020). Loop extrusion occurs in cells depleted for Pds5p or expressing mcd1p-Q266 (Costantino et al., 2020). While the pattern and the abundance of loops change in Pds5p depleted cells, they are largely unaffected in cells containing mcd1p-Q266 (Costantino et al., 2020). The ability of these mutants to loop extrude but not oligomerize, suggests that cohesin oligomers are not obligatory for loop extrusion *in vivo*. We suggest that cohesin oligomerization alters the inherent loop extruding activity of the monomers, perhaps by modulating its ability to promote symmetric or asymmetric looping (Davidson et al., 2019; Kim et al., 2019; Wang et al., 2017).

Budding yeast also regulates oligomer binding to chromosomes (this study). Our *trans*-biotinylation studies suggest that oligomers occur in S phase at all previously identified sites of cohesin binding (centromeres, pericentromeric regions, and the many CARS along chromosome arms). In mid-M, oligomers are specifically reduced at CARs (Figure 8A). This reduction in arm oligomers likely leads to a reduction in the robustness arm cohesion in mid-M, given the correlation between cohesin oligomers and cohesion maintenance (this study). Interestingly, the reduction of arm cohesion also occurs in mammalian prophase by Wpl1p-dependent inhibition of cohesin (Kueng et al., 2006). Thus, the reduction of arm cohesion is likely a conserved feature of mitosis (albeit by different mechanisms).

Proper mitotic chromosome segregation requires the establishment and maintenance of tension that results from the bipolar attachment of sister kinetochores and the surrounding cohesion (Makrantoni and Marston, 2018). This cohesion may be strengthened by the cohesin oligomers around the centromere, helping the tension to persist after bipolar attachment (Figure 8A). Similarly, bipolar attachment of homologs during meiosis I requires tension that is generated by cohesion distal to the crossover site. In *C*. *elegans*, COH-3/COH-4 cohesins are remodeled prior to anaphase 1 of meiosis persisting primarily distal to the crossover (Severson and Meyer, 2014). If COH-3/COH-4 forms oligomers, these oligomers may also serve to strengthen cohesion, facilitating its persistence under tension. Thus, these results may suggest that the regulation of oligomer persistence on the genome is a conserved feature of eukaryotes that is important in both meiosis and mitosis.

## Materials and Methods

### Yeast strains and media

Yeast strains used in this study are MATa cells with A364A background, and their genotypes are listed in Supplementary Table 1. Synthetic complete with low biotin (SC-Dex) was prepared by dissolving 1.4 g/L YNB-biotin (Sunrise Science Products), 1.6 g/L BSM powder (Sunrise Science Products), 5 g/L ammonium sulfate (Fisher Scientific), 20 g/L dextrose (Fisher Scientific) and 0.6 nM D-Biotin (Invitrogen) in distilled water and filter sterilized. Synthetic complete medium with raffinose (SC-Raff) for GAL induction was prepared similarly with SD-Dex, but 20 g/L dextrose was replaced with 20 g/L raffinose (Millipore-Sigma).

### Dilution plating assays

Cells were grown to saturation in YPD media at 30°C then plated in 10-fold serial dilutions on LEU- or FOA plates as described. Plates were incubated at 30°C for two days.

### Synchronous arrest in mid-M phase under auxin depletion conditions

#### G1 arrest

Asynchronous cultures of cells were grown in overnight culture to mid-log phase at 30°C in SC-Dex media (OD_600_ = 0.15). Cells were spun down and resuspended in fresh SC-Dex media containing 24 nM alpha factor (Millipore-Sigma). Cells were incubated at 30°C for 2 hours to induce arrest in G1 phase.

For depletion of AID-tagged proteins, auxin stock was prepared dissolving 44 mg auxin in 0.5 ml DMSO, and 0.05 ml auxin stock was added to each 25 ml medium (∼1 mM final). Cells were incubated for an additional 1 hour in alpha-Factor containing media.

#### Synchronous arrest in mid-M phase

0.1 mg/ml pronase (Millipore-Sigma, 10 mg/ml stock solution in water) was added to G1 arrested cells and the culture was incubated at 30°C for 10 minutes. The cells were spun down and resuspended in SC-Dex with 0.1 mg/ml pronase and 1 mM auxin. Nocodazole (Millipore-Sigma, 1.5 mg/ml in DMSO) was added to the culture dropwise to 12 μg/ml final, and ethyl acetate was added to a final concentration of 1.2%. Cells were incubated at 30°C for 2.5 hours to arrest in mid-M phase.

#### Biotinylation of Smc3p-AviTag reporters

Cells were harvested, resuspended in 0.8 ml SC-Dex with auxin and transferred to a low-binding Eppendorf tube. The biotinylation reaction was initiated by adding 10 nM D-biotin, incubated on a 30°C heat block for 7 minutes, and terminated by addition of 0.25 ml 80% TCA (Fisher).

### Protein extracts and western blotting

#### Total protein extracts

Cell equivalents of 2 OD_600_ were washed twice in cold PBS freshly supplemented with 0.5 mM PMSF (Millipore-Sigma) and resuspended in 0.3 ml lysis buffer (15% glycerol, 100 mM TRIS pH 8.0 and 0.2 mM PMSF). The cell lysate was prepared by bead beating on MP FastPrep 5G Homogenizer at top speed for 1 minute. The cell lysate was supplemented with 0.1 ml 4x SDS loading dye (16% SDS, 0.2% bromophenol blue and 20% β-mercaptoethanol) and heated at 95°C for ten minutes. The heated lysate was spun at a bench-top centrifuge at top speed for 3 minutes and the supernatant was stored in a freezer.

#### Streptavidin gel shifts

10 μl protein extract was mixed with 6 μl dilution buffer (1x PBS with 20% glycerol) and 2 μl streptavidin solution (Invitrogen, dissolved at 10 mg/ml in PBS), and incubated at room temperature for 10 minutes. The samples were loaded onto 8% SDS-PAGE gels, subjected to electrophoresis then transferred to PVDF membranes and analyzed by Western blot using standard laboratory techniques.

### Chromatin immunoprecipitation of biotinylated Smc3p

Cells treated with biotin pulse were fixed with 1% formaldehyde at room temperature for 1 hour and formaldehyde was quenched by incubating with 1 M glycine for 5 minutes. The cells were pelleted and washed 3x with FA-STD (50 mM HEPES pH 7.5, 150 mM NaCl, 1 mM EDTA pH 8.0, 1% Triton X-100, 0.1% SDS, 1% sodium deoxycholate and cOmplete protease inhibitors). The cells were suspended in 1 ml of FA-STD, beaten with glass beads twice on MP FastPrep 5G Homogenizer for 40 seconds. Chromatin was pelleted at 15,000 rpm for 15 minutes on a bench-top centrifuge, resuspended in 0.3 ml FA-STD, sheared on a Bioruptor Pico (Diagenode) for 10 min (30 s on/off cycling), and cleared by centrifugation at 15,000 rpm at 4°C for 10 minutes. Chromatin was then supplemented with additional 1% SDS and heated at 60°C for 5 minutes. 900 μl ice cold FA-STD was added to the tube to cool down the chromatin. The solution was supplemented with 1 mg/ml of BSA Cohn fraction V (Millipore-Sigma) and 0.1 mg/ml RNase A (Fisher), then incubated with 60 μl Dynabeads MyOne Streptavidin T1 at 4°C overnight. The beads were washed for 10 minutes at room temperature for each of the buffers listed below: (1) FA-STD with 1% SDS; (2)FA-STD with 3% SDS; (3) FA-STD with 3% SDS; (4) FA-STD with 1% SDS; (5) FA-STD with 1% SDS and 0.25 M LiCl; (6) FA-STD with 1% SDS and 0.5 M NaCl; (7) FA-STD with 1% SDS. The beads were then resuspended in 0.3 ml ChIP elution buffer (50 mM TRIS pH 8.0, 10 mM EDTA pH 8.0, 1% SDS and 0.5 mg/ml proteinase K) and incubated on a 65°C thermomixer overnight. The eluted DNA was purified with MinElute PCR purification kit (Qiagen) and analyzed by qPCR or NovaSeq (150bp paired-end, Illumina).

### Cohesin chromatin immunoprecipitation

Cohesin ChIP experiments were performed as described previously ((Eng et al., 2014; Robison et al., 2018)) with minor modifications. Cells were fixed with 1% formaldehyde at room temperature for 1 hour and formaldehyde was quenched by incubating with 1 M glycine for 5 minutes. The cells were spun down and washed 3x with FA-STD. Chromatin shearing was performed on a Bioruptor Pico (Diagenode) for 10 min (30 seconds on / 30 seconds off, 10 cycles). Immunoprecipitation was performed using polyclonal rabbit anti-Mcd1p (a gift from V. Guacci) antibodies and protein A Dynabeads (Invitrogen).

## Acknowledgements

We give special thanks to Gavin Schlissel for sharing the BirA plasmid and his guidance on chromatin immunoprecipitation of biotinylated proteins. We thank Vincent Guacci, Lorenzo Constantino, Zhouliang Yu and Kevin Boardman for inputs and discussions. This work used the Vincent J. Coates Genomics Sequencing Laboratory at UC Berkeley. This work was supported by the Helen Hay Whitney Foundation (S.X.) and the National Institutes of Health (grant 1R35 GM-118189-01 to D.E.K.).

