## Supplementary Figures and Tables for "Cohesin in space and time: architecture and oligomerization *in vivo*"

### Figure S1

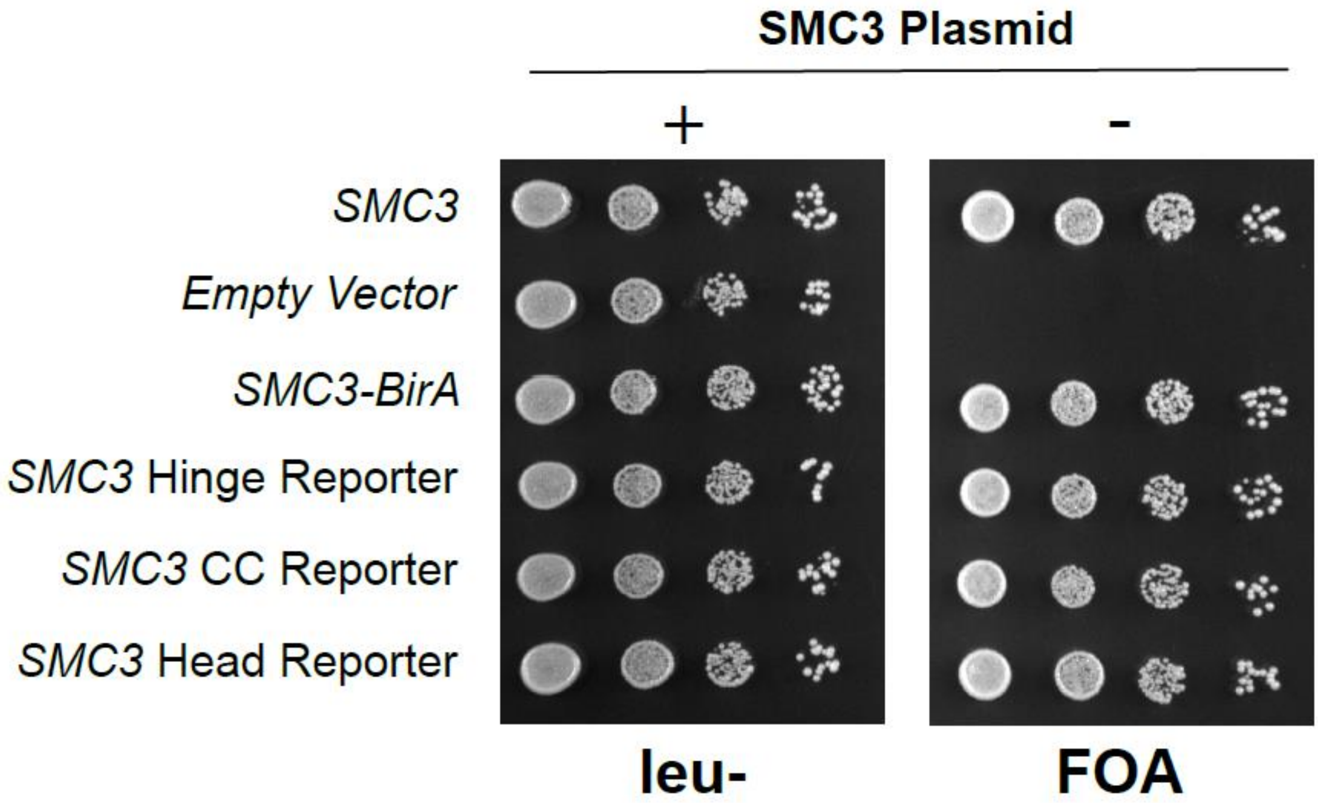


**Figure S1. Tagged SMC3 Alleles Support Cell Viability.**

Shuffle strains carrying CEN URA3 SMC3 plasmid and plasmid expressing the tagged SMC3 alleles. Strains were grown overnight in YPD medium and plated at 10-fold serial dilution on YPD or FOA plates.

### Figure S2

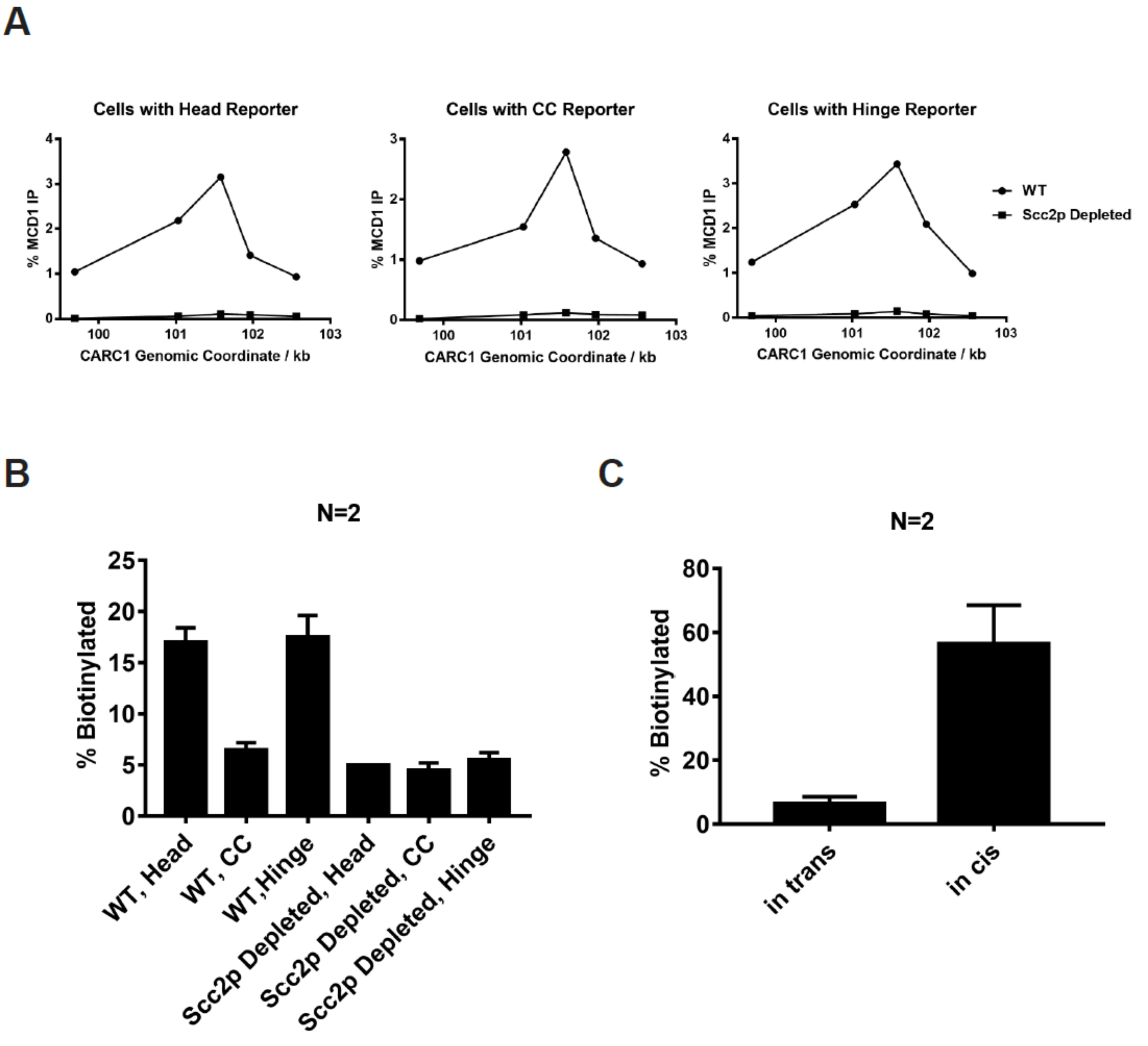


**Figure S2. Confirmation of SCC2-AID depletion and Quantitative Analysis of Figure 2.**

(A) Cohesin Failed to Bind Chromosome in scc2-depleted Cells. Cells from experiments described in Figure 2B were fixed for ChIP experiments. Cohesin bound chromosome was immunoprecipitated by anti-Mcd1 antibody and quantified by qPCR. Cohesin binding at a pericentric site (CARC1) in cells carrying wildtype or AID-tagged Scc2 was plotted against genomic coordinates.

(B) Quantitative analysis of Figure 2C. Ratios of Smc3p-AviTag reporter proteins biotinylated from two independent cultures were calculated based on band intensities (All error bars in this manuscript are indicative of standard deviations).

(C) Quantitative analysis of Figure 2E. Ratios of Smc3p-AviTag reporter proteins biotinylated from two independent cultures were calculated based on band intensities.

### Figure S3

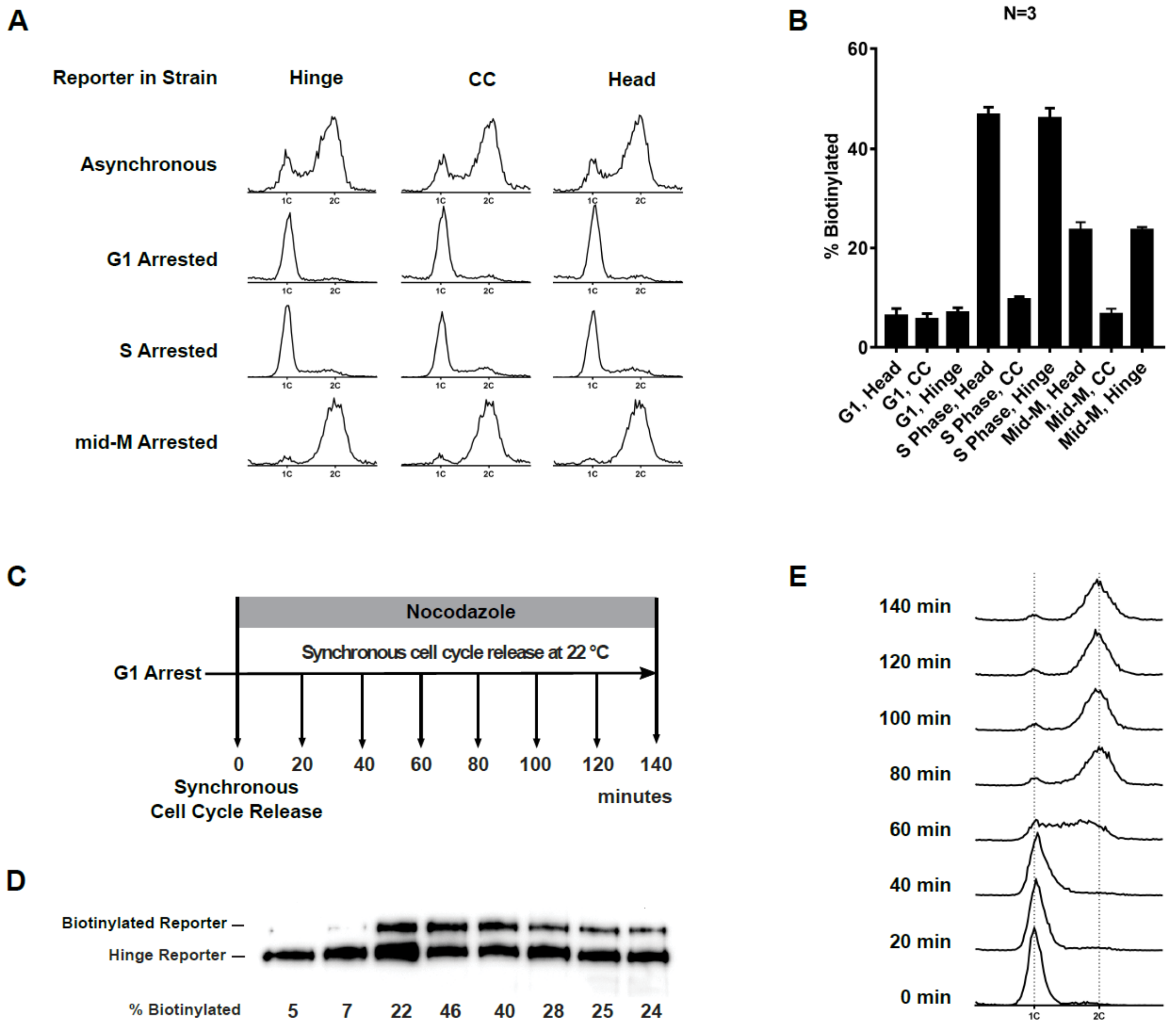


**Figure S3. Cohesin Oligomers in Cells Synchronously Released from G1 into mid-M Phase.**

(A) DNA content analysis in the G1, early S phase and mid-M phase arrested cells (left side, comparing with DNA content in asynchronous culture).

(B) Quantitative analysis of Figure 3C. Ratios of Smc3-AviTag reporter proteins biotinylated from three independent cultures were calculated based on band intensities.

(C) Regime used to assay cohesin oligomers in cells after synchronous G1 release. Cells carrying Smc3 hinge reporter were arrested in G1 and synchronously released into mid-M arrest. Aliquots of the culture were taken every 20 minutes and cohesin oligomerization levels in the cells were assayed with proximity biotinylation. Cells were preserved in 20% TCA and biotinylated reporters were assayed with streptavidin gel shift.

(D) Cohesin oligomerization levels during cell cycle progression. A basal level of cohesin oligomer was detected in G1 arrested cells. The ratio of cohesin in the oligomeric form increases during S Phase, and then drops to half of its peak level in mid-M arrested cells.

(E) Cell cycle progression was shown with DNA content analysis using flow cytometry.

### Figure S4

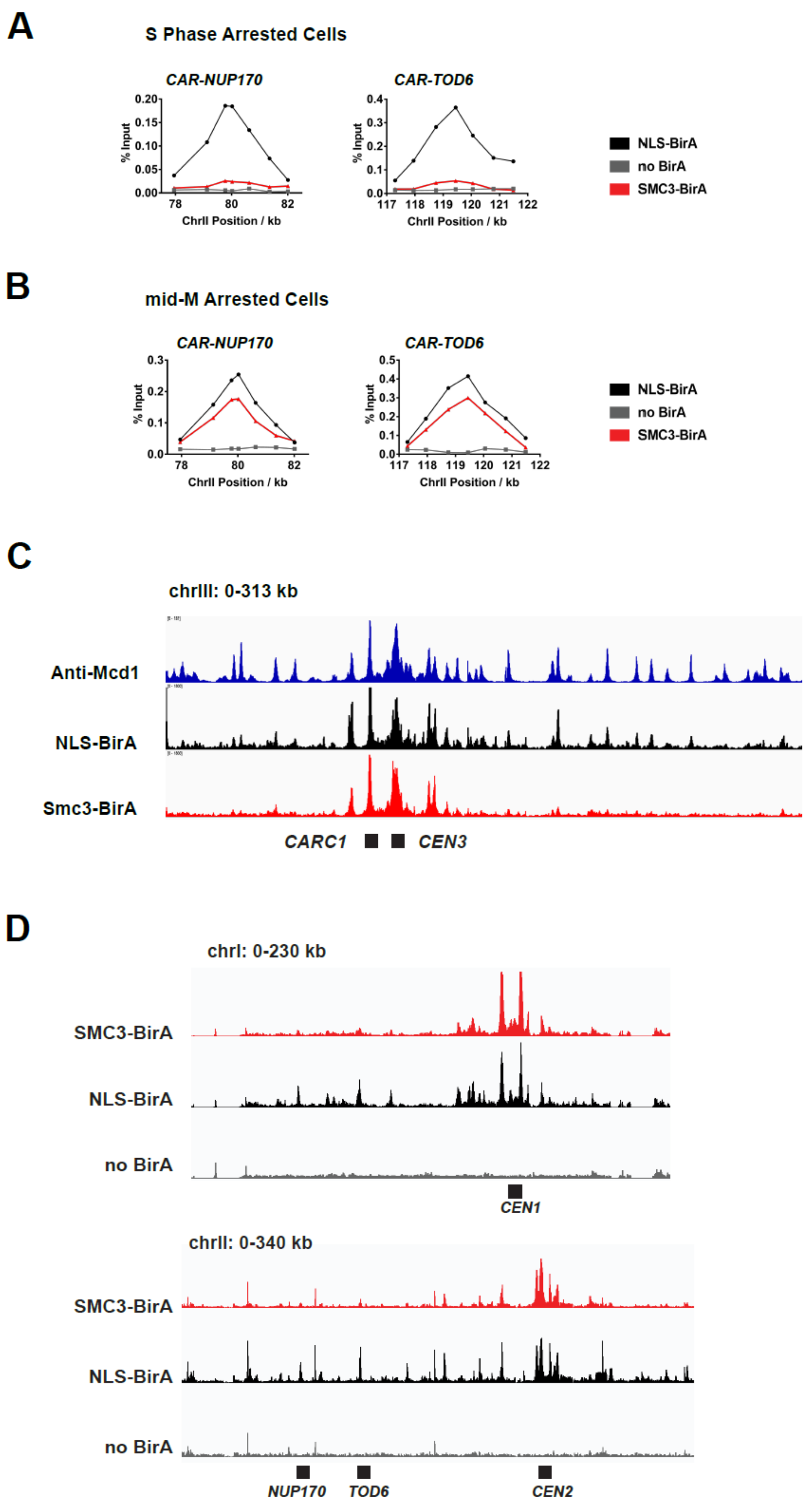
A
S Phase Arrested Cells
CAR-NUP170
CAR-TOD6
% Input
ChrII Position / kb
NLS-BirA
no BirA
SMC3-BirA
B
mid-M Arrested Cells
CAR-NUP170
CAR-TOD6
% Input
ChrII Position / kb
NLS-BirA
no BirA
SMC3-BirA
C
chrIII: 0-313 kb
Anti-Mcd1
NLS-BirA
Smc3-BirA
CARC1
CEN3
D
chrI: 0-230 kb
SMC3-BirA
NLS-BirA
no BirA
CEN1
chrII: 0-340 kb
SMC3-BirA
NLS-BirA
no BirA
NUP170
TOD6
CEN2

**Figure S4. ChIP Experiments Reveal the Centromeric and Pericentric Localization of Cohesin Oligomers.**

(A) Cohesin oligomers on the arm of chromosome II in S Phase arrested cells. Samples in Figure 4C were analyzed by qPCR using primers targeting CAR sites at the 3' end of NUP170 and TOD6. The two binding sites are both on the arm of chromosome II, as shown in (D).

(B) Cohesin oligomers on the arm of chromosome II in mid-M arrested cells. Samples in Figure 4D were analyzed by qPCR using primers targeting CAR sites at the 3' end of NUP170 and TOD6. The two binding sites are both on the arm of chromosome II, as shown in (D).

(C) Peaks of cohesin oligomers correlate with previously identified CAR site. Anti-Mcd1p ChIP-seq traces (Lorenzo *et al.* 2020) at chrIII was compared with ChIP-seq traces from Figure 4E.

(D) ChIP-seq traces showing localization of cohesin oligomers on chromosome I (top) and part of chromosome II (bottom). The figures were generated from the same sequencing data set used in Figure 4E. The trace with SMC3-BirA (red) shows position of cohesin oligomers; the trace with pGAL-BirA (black) shows cohesin binding sites; the no BirA trace (grey) is the negative control showing non-specific pull down from cells expressing no biotinylation enzyme.

### Figure S5

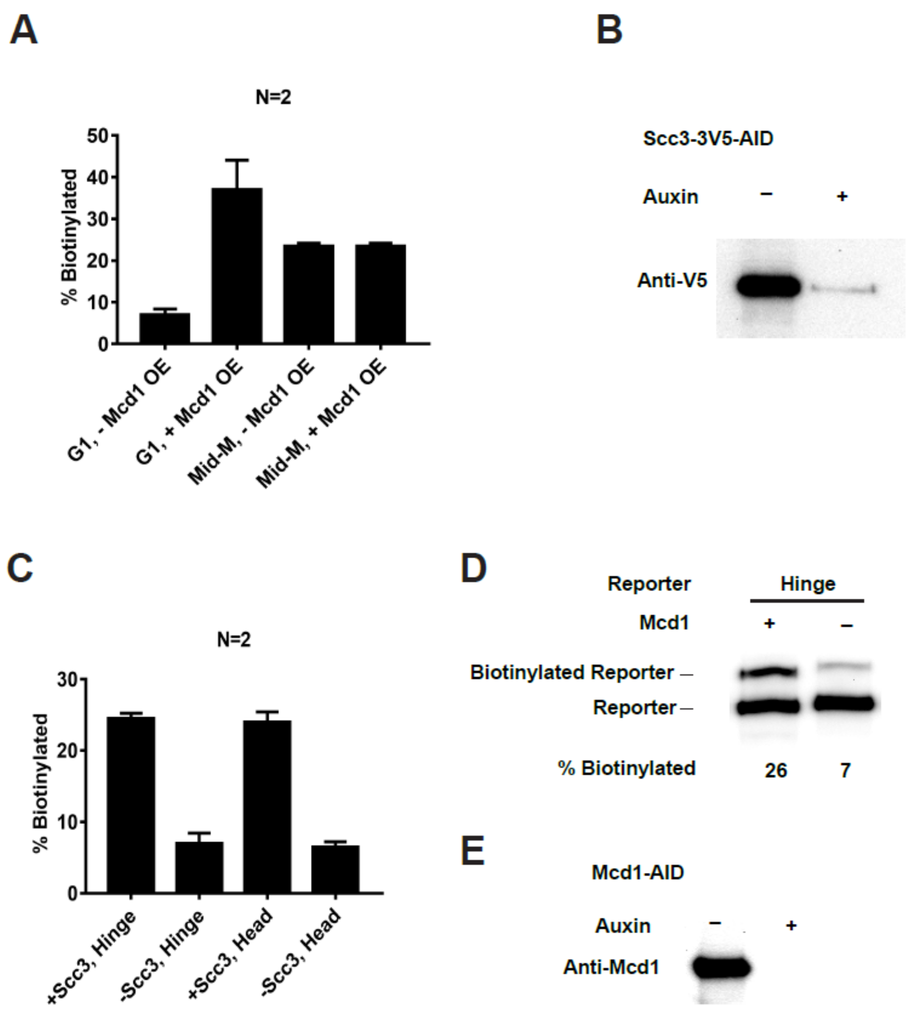


**Figure S5. Depletion of SCC2-AID and MCD1-AID.**

(A) Quantitative analysis of Figure 5B. Ratios of Smc3-AviTag reporter proteins biotinylated from independent cultures were calculated based on band intensities.

(B) Western blot confirms depletion of SCC3-AID.

(C) Quantitative analysis of Figure 5D. Ratios of Smc3-AviTag reporter proteins biotinylated from independent cultures were calculated based on band intensities.

(D) MCD1 is required for cohesin oligomerization. The plus signs indicate wild type cells expressing MCD1 while the minus signs indicate cells depleted of MCD1-AID. Cells were cultured following the experiment regime in Figure 5C. Cells were cultured in low biotin synthetic media, arrested in G1, and depletion of MCD1-AID was carried out by auxin addition. Synchronous cell cycle release was carried out in presence of auxin and cells were again arrested in mid-M phase by nocodazole. Proximity biotinylation experiments were carried out in the mid-M phase arrested cells.

(E) Confirmation of MCD1-AID depletion. Cells were cultured as (D) with (+) or without (-) adding auxin, and western blot was carried out with anti-Mcd1p antibody.

### Figure S6

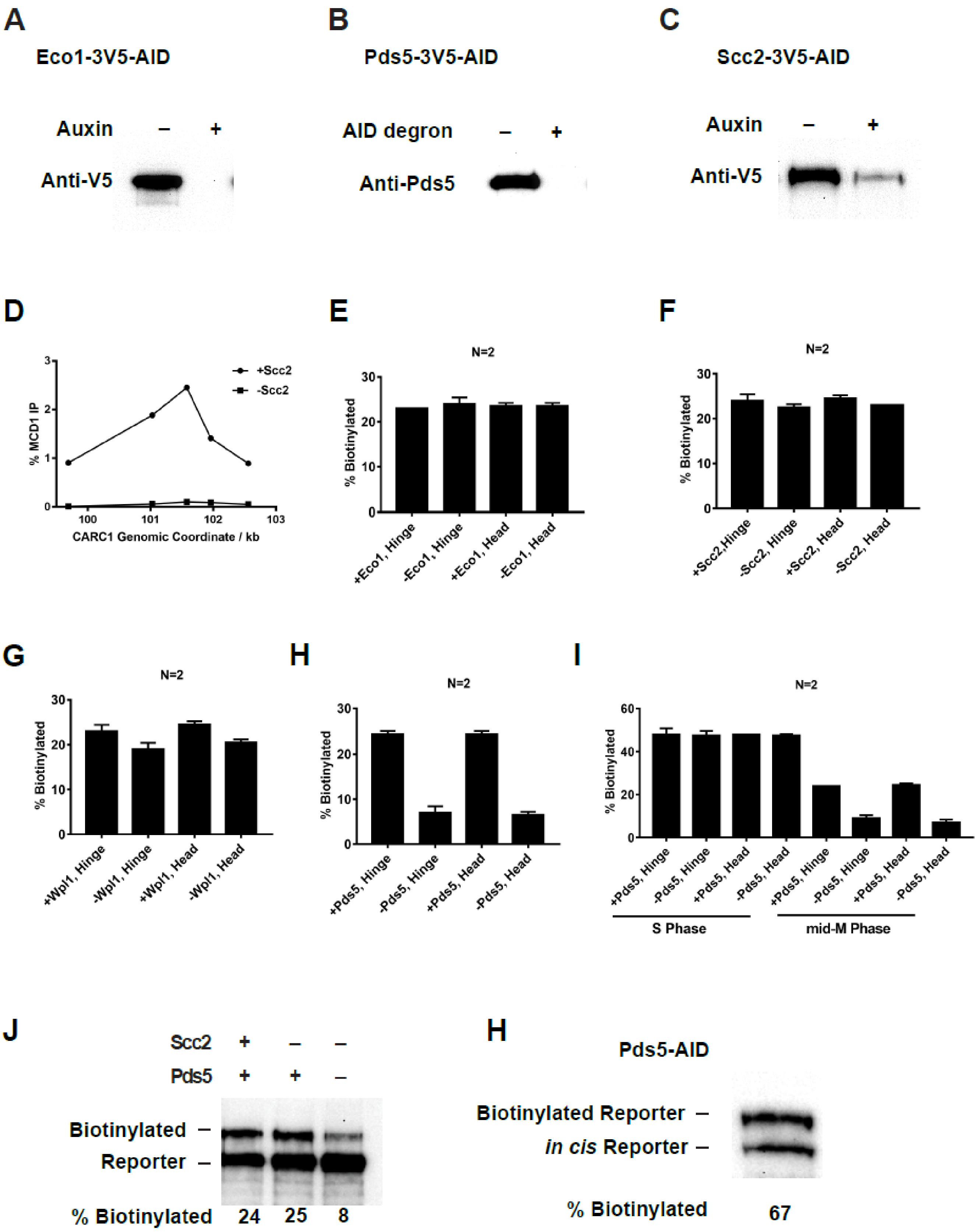
A
Eco1-3V5-AID
Auxin – +
Anti-V5
B
Pds5-3V5-AID
AID degron – +
Anti-Pds5
C
Scc2-3V5-AID
Auxin – +
Anti-V5
D
+Scc2
-Scc2
% MCD1 IP
CARC1 Genomic Coordinate / kb
E
N=2
% Biotinylated
+Eco1, Hinge
-Eco1, Hinge
+Eco1, Head
-Eco1, Head
F
N=2
% Biotinylated
+Scc2,Hinge
-Scc2, Hinge
+Scc2, Head
-Scc2, Head
G
N=2
% Biotinylated
+Wpl1, Hinge
-Wpl1, Hinge
+Wpl1, Head
-Wpl1, Head
H
N=2
% Biotinylated
+Pds5, Hinge
-Pds5, Hinge
+Pds5, Head
-Pds5, Head
I
N=2
% Biotinylated
+Pds5, Hinge
-Pds5, Hinge
+Pds5, Head
-Pds5, Head
S Phase
mid-M Phase
J
Scc2 + – –
Pds5 + + –
Biotinylated –
Reporter –
% Biotinylated 24 25 8
H
Pds5-AID
Biotinylated Reporter –
in cis Reporter –
% Biotinylated 67

**Figure S6. Depletion of Eco1, Pds5, Scc2 and Mcd1.**

(A) Western blot showing ECO1-AID depletion. Cells carrying hinge reporter were arrested in G1. The -auxin sample was collected before auxin addition, while the +auxin sample was collected one hour after adding auxin.

(B) Western blot showing PDS5-AID depletion. The same samples used for the first two lanes of Figure 6E were analyzed by western blotting using anti-Pds5p antibody. The first lane shows Pds5 protein level in cells with wild type Pds5. The second lane shows Pds5 protein level in cells with PDS5-AID.

(C) Western blot showing SCC2-AID depletion. Cells carrying hinge reporter were arrested in G1. The -auxin sample was collected before auxin addition, while the +auxin sample was collected one hour after adding auxin.

(D) Cohesin failed to bind chromosome in cells depleted of Scc2. Aliquots from cell cultures in the first two lanes of Figure 5C were fixed and subjected to ChIP against Mcd1.

(E) Quantitative analysis of Figure 6B. Ratios of Smc3-AviTag reporter proteins biotinylated from two independent cultures were calculated based on band intensities.

(F) Quantitative analysis of Figure 6C. Ratios of Smc3-AviTag reporter proteins biotinylated from two independent cultures were calculated based on band intensities.

(G) Quantitative analysis of Figure 6D. Ratios of Smc3-AviTag reporter proteins biotinylated from two independent cultures were calculated based on band intensities.

(H) Quantitative analysis of Figure 6E. Ratios of Smc3-AviTag reporter proteins biotinylated from two independent cultures were calculated based on band intensities.

(I) Quantitative analysis of Figure 6F. Ratios of Smc3-AviTag reporter proteins biotinylated from two independent cultures were calculated based on band intensities.

(J) Cohesin oligomer formed in mid-M arrested cells both on and off DNA are Pds5-dependent. Wild type cells (first lane), cells depleted of Scc2 (second lane) or cells depleted of both Scc2 and Pds5 (third lane) were synchronously release from G1 into mid-M phase and biotinylated. All three strains carry the Smc3p-BirA and the Smc3 hinge reporter.

(K) Fully assembled cohesin tetramers on DNA adopt the butterfly conformation. The strain carries an auxin-depletable PDS5-AID and a *cis*-biotinylation reporter allele (as shown in Figure 2D, right) of SMC3. The strain was arrested in mid-M using nocodazole and auxin was added to the culture to deplete Pds5p-AID. The cells were then treated with biotin pulse. *Trans*-biotinylation was abolished in this condition (Figure 6E) and the biotinylation shown is the result of intramolecular reaction.

### Figure S7

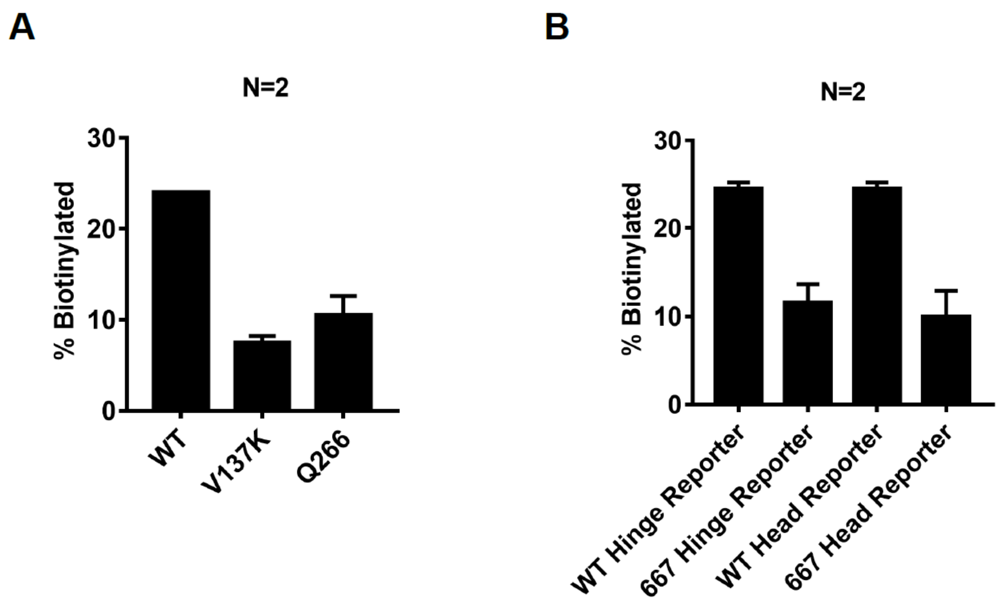


Figure S7. Quantification of Smc3 Reporter Biotinylation in Cells Expressing mcd1-V137K, mcd1-Q266 or smc3-D667.

(A) Quantitative analysis of Figure 7B. Ratios of Smc3-AviTag reporter proteins biotinylated from two independent cultures were calculated based on band intensities.

(B) Quantitative analysis of Figure 7D. Ratios of Smc3-AviTag reporter proteins biotinylated from two independent cultures were calculated based on band intensities.

**Table S1. Key Reagents**

| Reagent Name | Source | Identifier | Additional Information |
|---|---|---|---|
| Genetic reagent (*S. cerevisiae* ) | This paper | | Table S2 |
| qPCR Primers | IDT | | Table S3 |
| Rabbit polyconal anti-Mcd1 | D. Koshland via Covance | Anti-Mcd1 (555) | WB (1:10,000) ChIP (1:1,000) |
| Rabbit polyconal anti-Pds5 | D. Koshland via Covance | Anti-Pds5 (556) | WB (1:20,000) |
| Mouse monoclonal Anti-HA (12CA5) | Roche | 11667203001 | WB (1:8,000) |
| Mouse monoclonal Anti-V5 | Invitrogen | 46-0705 | WB (1:8,000) |
| Goat polyclonal HRP Anti-Rabbit | Biorad | 170-6515 | WB (1:8,000) |
| Goat polyclonal HRP Anti-Mouse | Biorad | 170-6516 | WB (1:5,000) |
| Rabbit polyclonal Anti-Tub2p | P.Meluh via Covance | | WB (1:20,000) |
| Protein A Dynabeads | Invitrogen | 10002D | |
| Dynabeads MyOne Streptavidin T1 | Invitrogen | 65601 | |
| Streptavidin | Invitrogen | S888 | |
| Auxin | Millipore-Sigma | C9911 | 1 mM final concentration, 0.5M stock in DMSO |
| Alpha Factor | Millipore-Sigma | T6901 | 24 nM final concentration, 10 nM stock |
| Hydroxyurea | Millipore-Sigma | H8627 | 0.2 M final concentration |
| Nacodazole | Millipore-Sigma | M1404 | 0.012 mg/ml final concentration, 1.5 mg/ml stock in DMSO |
| Ethyl acetate ACS | Millipore-Sigma | 319902 | |
| Pronase protease | Millipore-Sigma | 537088 | 10 mg/ml stock in water |
| YNB-biotin | Sunrise Science Products | 1523-100 | |
| BSM powder | Sunrise Science Products | 1387-100 | |
| D-Biotin | Invitrogen | B20656 | |

**Table S2. Yeast Strains**

| | |
|---|---|
| **VG3620-4C** | TIR1-CaTRP1 LacO-NAT::lys4 GFPLacI-HIS3:his3-11,15 leu2-3,112 ura3-52 bar1 GAL+ |
| **SX48B** | TIR1-CaTRP1 GFPLacI-HIS3:his3-11,15 LacO-NAT::lys4 SMC3-A1089-V5-BirA ura3-52 bar1 GAL+ SMC3-(P533-AviTag-6HA)-LEU2:leu2-3,112 |
| **SX48E** | TIR1-CaTRP1 GFPLacI-HIS3:his3-11,15 LacO-NAT::lys4 SMC3-A1089-V5-BirA ura3-52 bar1 GAL+ SMC3-(V966-AviTag-6HA)-LEU2:leu2-3,112 |
| **SX48F** | TIR1-CaTRP1 GFPLacI-HIS3:his3-11,15 LacO-NAT::lys4 SMC3-A1089-V5-BirA ura3-52 bar1 GAL+ SMC3-(A1089-AviTag-6HA)-LEU2:leu2-3,112 |
| **SX49B** | TIR1-CaTRP1 GFPLacI-HIS3:his3-11,15 LacO-NAT::lys4 SMC3-A1089-V5-BirA ura3-52 bar1 GAL+ smc3-D667-(P533-AviTag-6HA)-LEU2:leu2-3,112 |
| **SX49F** | TIR1-CaTRP1 GFPLacI-HIS3:his3-11,15 LacO-NAT::lys4 SMC3-A1089-V5-BirA ura3-52 bar1 GAL+ smc3-D667-(A1089-AviTag-6HA)-LEU2:leu2-3,112 |
| **SX66B** | TIR1-CaTRP1 GFPLacI-HIS3:his3-11,15 LacO-NAT::lys4 SMC3-A1089-V5-BirA ura3-52 bar1 GAL+ SMC3-(P533-AviTag-6HA)-LEU2:leu2-3,112 wplΔ::HPH |
| **SX66F** | TIR1-CaTRP1 GFPLacI-HIS3:his3-11,15 LacO-NAT::lys4 SMC3-A1089-V5-BirA ura3-52 bar1 GAL+ SMC3-(A1089-AviTag-6HA)-LEU2:leu2-3,112 wplΔ::HPH |
| **SX73** | TIR1-CaTRP1 LacO-NAT::lys4 GFPLacI-HIS3:his3-11,15 SMC3-(A1089-AviTag-6HA)-LEU2:leu2-3,112 ura3-52 bar1 GAL+ |
| **SX74** | TIR1-CaTRP1 GFPLacI-HIS3:his3-11,15 SMC3-A1089-V5-BirA ura3-52 bar1 GAL+ SMC3-(A1089-dead-AviTag-6HA)-LEU2:leu2-3,112 |
| **SX80B** | TIR1-CaTRP1 LacO-NAT::lys4 GFPLacI-HIS3:his3-11,15 GAL1p-NLS-V5-BirA-URA3::ura3-52 SMC3-(P533-AviTag-6HA)-LEU2:leu2-3,112 leu2-3,112 ura3-52 bar1 GAL+ |
| **SX80E** | TIR1-CaTRP1 LacO-NAT::lys4 GFPLacI-HIS3:his3-11,15 GAL1p-NLS-V5-BirA-URA3::ura3-52 SMC3-(V966-AviTag-6HA)-LEU2:leu2-3,112 leu2-3,112 ura3-52 bar1 GAL+ |

| | |
|---|---|
| **SX80F** | TIR1-CaTRP1 LacO-NAT::lys4 GFPLacI-HIS3:his3-11,15<br>GAL1p-NLS-V5-BirA-URA3::ura3-52<br>SMC3-(A1089-AviTag-6HA)-LEU2:leu2-3,112<br>leu2-3,112 ura3-52 bar1 GAL+ |
| **SX81B** | TIR1-CaTRP1 GFPLacI-HIS3:his3-11,15 LacO-NAT::lys4<br>SMC3-A1089-V5-BirA ura3-52 bar1 GAL+<br>SMC3-(P533-AviTag-6HA)-LEU2:leu2-3,112<br>ECO1-3V5-AID2:G418 |
| **SX81F** | TIR1-CaTRP1 GFPLacI-HIS3:his3-11,15 LacO-NAT::lys4<br>SMC3-A1089-V5-BirA ura3-52 bar1 GAL+<br>SMC3-(A1089-AviTag-6HA)-LEU2:leu2-3,112<br>ECO1-3V5-AID2:G418 |
| **SX82B** | TIR1-CaTRP1 GFPLacI-HIS3:his3-11,15 LacO-NAT::lys4<br>SMC3-A1089-V5-BirA ura3-52 bar1 GAL+<br>SMC3-(P533-AviTag-6HA)-LEU2:leu2-3,112<br>SCC2-3XFlag-AID2:HYG |
| **SX82F** | TIR1-CaTRP1 GFPLacI-HIS3:his3-11,15 LacO-NAT::lys4<br>SMC3-A1089-V5-BirA ura3-52 bar1 GAL+<br>SMC3-(A1089-AviTag-6HA)-LEU2:leu2-3,112<br>SCC2-3XFlag-AID2:HYG |
| **SX115** | trp1-1::G418-TIR1 GFPLacI-HIS3:his3-11,15 LacO-NAT::lys4<br>SMC3-(P533-AviTag-6HA)-(A1089-V5-BirA)<br>leu2-3,112 ura3-52 bar1 GAL+ |
| **SX122B** | TIR1-CaTRP1 GFPLacI-HIS3:his3-11,15 LacO-NAT::lys4<br>SMC3-A1089-V5-BirA ura3-52 bar1 GAL+<br>SMC3-(P533-AviTag-6HA)-LEU2:leu2-3,112<br>PDS5-3V5-AID2:G418 |
| **SX122F** | TIR1-CaTRP1 GFPLacI-HIS3:his3-11,15 LacO-NAT::lys4<br>SMC3-A1089-V5-BirA ura3-52 bar1 GAL+<br>SMC3-(A1089-AviTag-6HA)-LEU2:leu2-3,112<br>PDS5-3V5-AID2:G418 |
| **SX129** | trp1-1::cgTRP1-TIR1 GFPLacI-HIS3:his3-11,15 bar1 GAL+<br>MCD1-AID-G418 (no tag) ura3-52::mcd1-Q266-6HA-URA3<br>SMC3-A1089-V5-BirA SMC3-(P533-AviTag-6HA)-LEU2:leu2-3,112 |
| **SX130** | trp1-1::cgTRP1-TIR1 GFPLacI-HIS3:his3-11,15 bar1 GAL+<br>MCD1-AID-G418 (no tag) ura3-52::mcd1-V137K-6HA-URA3<br>SMC3-A1089-V5-BirA SMC3-(P533-AviTag-6HA)-LEU2:leu2-3,112 |
| **SX134** | trp1-1::cgTRP1-TIR1 GFPLacI-HIS3:his3-11,15 bar1 GAL+<br>MCD1-AID-G418 (no tag) ura3-52::MCD1-6HA-URA3<br>SMC3-A1089-V5-BirA SMC3-(P533-AviTag-6HA)-LEU2:leu2-3,112 |
| **SX136B** | TIR1-cgTRP1:trp1-1<br>CARC1::4x(tetO-lacO) his3-11,15:HUalpha-BirA-HIS3<br>leu2-3,112:SMC3-(P533-AviTag-6HA)-LEU2<br>ura3-52 lys2-801 bar1 GAL+ |

| | |
|---|---|
| **SX136E** | TIR1-cgTRP1:trp1-1<br>CARC1::4x(tetO-lacO) his3-11,15:HUalpha-BirA-HIS3<br>leu2-3,112:SMC3-(V966-AviTag-6HA)-LEU2<br>ura3-52 lys2-801 bar1 GAL+ |
| **SX136B** | TIR1-cgTRP1:trp1-1<br>CARC1::4x(tetO-lacO) his3-11,15:HUalpha-BirA-HIS3<br>leu2-3,112:SMC3-(A1089-AviTag-6HA)-LEU2<br>ura3-52 lys2-801 bar1 GAL+ |
| **SX155** | TIR1-CaTRP1 GFPLacI-HIS3:his3-11,15 LacO-NAT::lys4<br>SMC3-A1089-V5-BirA ura3-52 bar1 GAL+<br>SMC3wt-13myc-AviTag-LEU2:leu2-3,112 |
| **SX158** | TIR1-CaTRP1 LacO-NAT::lys4 GFPLacI-HIS3:his3-11,15<br>SMC3-A1089-V5-BirA PDS5-3V5-AID2:G418<br>scc2-3xFLAG-AID2:HYG leu2-3,112::SMC3-P533-Avi6HA-LEU2<br>ura3-52 bar1 GAL+ |
| **SX172** | TIR1-CaTRP1 GFPLacI-HIS3:his3-11,15 LacO-NAT::lys4<br>SMC3-13myc-AviTag-LEU2:leu2-3,112<br>ura3-52 bar1 GAL+ |
| **SX173** | TIR1-CaTRP1 LacO-NAT::lys4 GFPLacI-HIS3:his3-11,15<br>GAL1p-NLS-V5-BirA-URA3::ura3-52 ura3-52 bar1 GAL+<br>SMC3-13myc-AviTag-LEU2:leu2-3,112 |
| **SX220B** | TIR1-CaTRP1 LacO-NAT::lys4 GFPLacI-HIS3:his3-11,15<br>SMC3-A1089-V5-BirA leu2-3,112:SMC3-(P533-AviTag-6HA)-LEU2 |
| **SX220F** | TIR1-CaTRP1 LacO-NAT::lys4 GFPLacI-HIS3:his3-11,15<br>SMC3-A1089-V5-BirA leu2-3,112:SMC3-(A1089-AviTag-6HA)-LEU2<br>scc3-3V5-AID2-G418 ura3-52 bar1 GAL+ |
| **SX221B** | TIR1-cgTRP1:trp1-1 ura3-52 lys2-801 bar1 GAL+<br>CARC1::4x(tetO-lacO) his3-11,15:HUalpha-BirA-HIS3<br>scc2-3xFLAG-AID2:HYG<br>leu2-3,112:SMC3-(P533-AviTag-6HA)-LEU2 |
| **SX221E** | TIR1-cgTRP1:trp1-1 ura3-52 lys2-801 bar1 GAL+<br>CARC1::4x(tetO-lacO) his3-11,15:HUalpha-BirA-HIS3<br>scc2-3xFLAG-AID2:HYG<br>leu2-3,112:SMC3-(V966-AviTag-6HA)-LEU2 |
| **SX221F** | TIR1-cgTRP1:trp1-1 ura3-52 lys2-801 bar1 GAL+<br>CARC1::4x(tetO-lacO) his3-11,15:HUalpha-BirA-HIS3<br>scc2-3xFLAG-AID2:HYG<br>leu2-3,112:SMC3-(A1089-AviTag-6HA)-LEU2 |
| **SX222** | TIR1-CaTRP1 LacO-NAT::lys4 GFPLacI-HIS3:his3-11,15<br>pGAL1-(NLS-BirA)-URA3::ura3-52<br>SMC3-A1089-(dead-AviTag-6HA)-LEU2::leu2-3,112 bar1 GAL+ |

**Table S3. Primers used for chromatin Immunoprecipitation (ChIP)**

##### *Arm TRM1 CAR*

VG641/VG642 (AAAGAAGCAGGGGTAGAGAAGC / ATCAGCAGCGGTGATTACAC)
VG625/VG626 (CCAGCCAGATATTATGGGCAAG / TCCTAGACCTGGTGGAAAAAGC)
VG627/VG628 (CTTATAGTTCCCAAGGCATCCC / CCAAACTCGTTGTTCTCGATCC)
VG629/VG630 (TCTTCGTGCGCGAGGATATG / CGAACATTTCCGGACAATTGC)
VG633/VG634 (CCAATCGTATAACGGAGCATTGG / TGGTGCCAGAAGATATCAACG)
VG637/VG638 (GCGCGATACCATTCAGAACATC / TTAAAGTGGGCCCAAGACCAG)
VG639/VG640 (GGGCATCACCTTTTCGTAAGC / TGATCCACCTGTCATTTCGC)
VG643/VG644 (ACCCTTCTGTTCCAGTTTGC / GTTGCCTCCGGAGCAAATTC)

##### *Arm NUP170 CAR*

NUP170-1F/1R (CAATTCTAACGGCAGGGTTATTG / CGACTTCGCTTTCCTCGTATAA)
NUP170-3F/3R (GCATGTGAAGTTGCTGGTATTC / ACAGCTCACTTGTGCTCAATA)
NUP170-4F/4R (CGTTAAGAACAGCGGCAATAAT / ACACGTACATTACCCTGCTATC)
NUP170-4aF/4aR (CGAACAGGTCAGAGAGA / ACAGTAAGGTGGAGTTAATG)
NUP170-5F/5R (GTAAGGTCAGCAGGAACTAGATATT / GCTGACAGGTTCAAGAATAGGA)
NUP170-6F/6R (TTCTGTTGACCAGCTCTCTTG / GCTATTAGAGGCTTGCCAGAA)
NUP170-7F/7R (GGCACGGACAAATGTGAATAAG / TCGTCGTTTGAAGGGTGAAA)

##### *Arm TOD6 CAR*

TOD6-1F/1R (GGCCTATCTTGTCTCATCATCTT / ATCGCATCGCATCTCATCTC)
TOD6-2F/2R (TTGGGCTGGAAGGAGATTG / TCGCATACACGCCGAATAG)
TOD6-3F/3R (CAAGACAGACCACGCAAATG / CTCTAGACCACGGGTGTTTATT)
TOD6-4F/4R (ATACCCATCGCCGCTTATTC / GATGATGATGAGGATGGGAAGAG)
TOD6-5F/5R (TGCTCACTACTTCTTCTTCTTCTT / ACTTAGAACCCACCGCATTAG)
TOD6-6F/6R (CTCTGTATCCTCTCTCTCCGTTAG / GGCGGGAGAATGTCTTGTATT)
TOD6-7F/7R (GATTAGAGGACCGGATGATGTT / CAGCGAGGAGTTGAAGGTTAG)

##### *Pericentric CARC1*

TE367/TE368 (AAAGGTGCCCCAAGAAAAGG / AGCACTTTACTCGCTTGTGG)
TE310/TE311 (TAAAGCATTGACGCCAGAGC / AAGTACGCGTACGAAGCATC)
TE308/TE309 (TCCTGGAATGGAGACCGTTTTC / AGCCGACAAATTTCGTGCAC)
TE373/TE374 (ACTTTGGTTTTCCGGTGTGC / CCAGCGATGAGATGCGAAAAG)
TE377/TE378 (TCGCTTTTCGCATCTCATCG / AGCGGGCGGGTTATAAATAAC)
TE533/TE534 (ACCTTCTACTTCCATGCCGTTG / TGCGTGCCGATGTAGAATTG)

#### *CEN* primers

##### *CEN4* flanking primers

BR463/BR464 (CATGATTCGCCGGGTAAATA / GCACTAGCCAATTTAGCACTTC)

BR465/BR466 (AAAATGCCGAGGCTTTCATA / TGACGATAAAACCGGAAGGA)

##### *CEN14* flanking primers

TE442/TE443 (TTAAAGCGGCTGAGTATGGC / TTTCCTCCATTGCTCTCTACGG)

TE446/TE447 (ACTAAAAGTGCCCCAAACGG / AGGAGCAGGGTAGCATAAACC)
